# Description of three new predator species of Litostomatea (Alveolata, Ciliophora), including sequencing and annotation of their mitochondrial genomes

**DOI:** 10.1101/2025.07.17.664916

**Authors:** Alessandro Allievi, Leandro Gammuto, Wanying Liao, Sergei I. Fokin, Giulio Petroni, Letizia Modeo, Valentina Serra

## Abstract

Three new species of haptorian ciliates *Lacrymaria venatrix* sp. nov., *Phialina famelica* sp. nov., and *Chanaea vermicularis* sp. nov., recovered from water bodies of Tuscany, Italy, are described according to the standards of the Next Generation Taxonomy, which include morphology, ultrastructure, phylogeny, and associated microbial consortium analysis plus mitochondrial genome sequencing and annotation. *Lacrymaria venatrix* is a species with a small spindle-shaped trunk, a highly contractile neck that equals the trunk in length at its full extension, and a macronucleus consisting of two nodules; it is mostly similar to *L. songi* but diverges in the denser ciliature, the closer macronuclear nodules, and the single micronucleus. *Phialina famelica* has a blunter spindle-shaped trunk with a shallow oral bulge and an elongated macronucleus; it is mostly similar to *P. salinarum* and *P. serranoi*, but differs in the freshwater habitat, and by having fewer but denser kineties. *Chaenea vermicularis* differs from its congenerics by a modest contractility combined with a greater elongation ability, the mucocysts organized into multiple rows forming stripes, and the several hundreds of macronuclear nodules. The 18S rDNA-based phylogenetic analysis supports previous observations of the paraphyly of the genera *Lacrymaria* and *Phialina*, that need a deep systematics revision. The genus *Chaenea* seems not to be challenged, though its position within Haptoria remains not fully clear, and *C. vermicularis* sp. nov. appears rather divergent from its congeners.

## INTRODUCTION

The class Litostomatea Small and Lynn 1981 comprises ciliated organisms (Phylum Ciliophora) that are aerobic predators (subclasses Rhynchostomatia and Haptoria), or that inhabit the gut of both invertebrate and vertebrate hosts, as anaerobic endosymbionts (subclass Trichostomatia) (Cedrola et al., 2020; Foissner and Xu, 2007). The subclass Haptoria Corliss 1974 is distinguished by other Litostomatea by the presence of toxicysts around the oral region, which is surrounded by a ring of dikinetids, by a denser patch of somatic cilia known as the dorsal brush (with the somatic ciliature being otherwise uniform), and by various features of cortex ultrastructure (Lynn, 2008; Vďačný et al., 2014). Haptorians are specialized predators of other unicellular organisms, particularly ciliates of similar size, using their toxicysts either to immobilize or kill their preys. Examples include genera such as *Litonotus*, *Loxophyllum*, *Spathidium*, *Didinium*, *Chaenea*, and the *Lacrymaria* + *Phialina* species cluster. The systematic of the subclass Haptoria, has been highly debated in the previous years, and recent phylogenetic analyses based on molecular data even showed it to be paraphyletic, with subclass Trichostomatia nesting within family Spathidiidae (Vďačný et al. 2014, Huang et al. 2018).

In this context, the ciliate genera *Lacrymaria* Bory, 1824 and *Phialina* Bory, 1824 have both been described very early in the history of protistology (Müller, 1773, 1786; Bory de Saint-Vincent, 1824; Ehrenberg, 1831). They have had a rather complex taxonomic history, with the diagnostic features discriminating between the two (first mouth position, then “neck” length), and even the status of *Phialina* as separate genus, have changed multiple times. Both genera have a body divided into a spindle- or bottle-shaped “trunk”, with a highly variable width-to-length ratio and a number of longitudinal or spiral kineties, a distinct “head” covered by a separate set of kineties and often containing the toxicysts, and a dome- or cone-shaped “oral bulge” surrounding the mouth. The anterior portion of the trunk may extend into a highly contractile and flexible neck somewhat narrower than the head. Since Foissner (1983) the presence of the neck has been considered the discriminating trait between *Lacrymaria* and *Phialina*. Unlike the head, the neck is not sharply separated from the trunk. In this trait they both differ from the similar genera *Trachelophyllum* and *Enchelys*, which have no distinct head, and *Lagynus*, which has multiple anterior segments (see Foissner, 1983, Foissner et al., 1995). Somatic kineties wrap around the whole length of the trunk at an angle from the body axis. The nuclear apparatus is variable in shape and number, being, indeed, a feature very useful to differentiate among the species, most frequently it consists in one or two ellipsoidal nodules. In general, many species of *Lacrymaria* and *Phialina* have been poorly described over the years. Some have been observed only once, and for only a few of them (e.g., *L. marina*, *L. maurea*, *L. olor*, *P. caudata*, *P. clampi*, *P. pupula*, *P. salinarum*, *P. vermicularis*, *P. vertens*) taxonomically relevant molecular marker sequences have been deposited in online databases.

The genus *Chaenea* Quennerstedt, 1867 is another member of subclass Haptoria. During the latest decades, some new representatives of freshwater or (mainly) marine ciliates belonging to the genus were found and described with some references to older literature (Carey, 1992; Foissner et al., 1995; Petz et al., 1995; Song and Packroff, 1997; Jankowski, 2007; Gao et al., 2008; Song et al., 2009; Zhang et al., 2012; Kwon et al., 2014; Fan et al., 2015; Jiang et al., 2021). Even a new family Chaeneidae within the order Lacrymariida Lipscomb and Riordan, 1990 was proposed for them a decade ago (Kwon et al., 2014). The type species of the genus is *C. vorax* Quennerstedt, 1867, a ciliate predator frequently found in marine littoral environments.

Ciliates of the genus *Cheanea* are narrow, relatively big in length (up to ∼ 500 µm *in vivo*), often worm-like or serpentine in shape with a weakly contractile body and a limited number (8-20) of somatic kineties. A small rounded oral bulge, located on the anterior region shaped like a head, is always present with rod-shaped toxicysts inserted, as well as a dorsal brush consisting of four ciliary rows with a peculiar organization. The macronucleus usually consists of nodules, more or less numerous and differently distributed in the cell. In the literature, about 20 species are known, but some of them should be rechecked to confirm their validity considering that, although many are morphologically distinct taxa and a key to the identification of the species has been recently offered (Fan et al., 2015), several species have been reported only once, and sometimes only living observations have been provided (Fan et al., 2015). Additionally, molecular investigation (Kwon et al., 2014) showed a very high similarity between some of the few species for which gene data are available, i.e., *C. vorax*, *C. teres*, and *C. mirabilis* (99.2 – 100%; a difference of only 0-9 nucleotides). Due to the lack of detailed morphological data on the sequenced C*haenea* populations (especially for *C. teres*), Kwon et al. (2014) suggested that *C. teres* EF486860 and *C. vorax* DQ190461 have been possibly misidentified. Thus, currently, both morphological and molecular classifications of *Chaenea* representatives are partly controversial, and Vd’ačny et al. (2014) indicated the genus as *incertae sedis* within the Haptoria. However, given that the family Chaeneidae has been identified as the deepest branch within the Litostomatea, its possible elevation to the order rank has recently been proposed by Zhang et al. (2024) as visible in Figures 6 and 7 of their paper, although this suggestion has not been formally discussed.

In the context of such complex and mutable systematic framework, the introduction of new standards of taxonomic description, combining multiple techniques and areas of interest, may be useful to stabilize taxa for which phylogenetic relationships have undergone severe changes during time (even reverting to previous positions multiple times), as well as establish new, reliable ones.

Next Generation Taxonomy, or “NGTax” (Serra et al., 2020), is a recently established, continuously updated taxonomic workflow which provides such a standard. NGTax combines morphological, molecular, and phylogenetic/phylogenomic considerations in order to give a rounded, multi-dimensional description of an organism including all traits that may be of taxonomic interest (e.g., traditional morphology, ultrastructure, associated microbiome description, different nuclear molecular markers or mitochondrial genome). It has been successfully applied on ciliates scale (Serra et al., 2020; Liao et al., 2021a; Liao et al., 2021b; Serra et al., 2021; Fokin et al., 2023; Xu et al., 2025) as well as on metazoans (Gammuto et al., 2024).

In the present study, we describe three species of the Class Litostomatea, namely *Lacrymaria venatrix*, *Phialina famelica*, and *Chaenea vermicularis*, following the multidisciplinary NGTax approach. We present a large set of data for each species including morphological, morphometric, ultrastructural, and molecular information, such as an 18S rDNA-based phylogeny, analyses on the mitochondrial genome (mitogenome) sequence and synteny, to provide their best characterization and allow their comparison. Previous research suggested the genera *Lacrymaria* and *Phialina* need a thorough systematics revision due to their paraphyly, and we discuss this issue in detail. Additionally, we investigated the microbial consortium associated with the three predatory species using molecular analysis, fluorescence in situ hybridization (FISH) and transmission electron microscopy.

## MATERIALS AND METHODS

### Sample collection and culture setting

The three novel species were collected from three different localities: *Lacrymaria venatrix* sp. nov. (strain PDF5b_Lac3), from Padule di Fucecchio (43.807°N, 10.810 °E), a wetland environment in the province of Pistoia (Tuscany, Italy). *Phialina famelica* sp. nov. (strain I4) was collected from. A wastewater treatment plant (WWTP) managed by Cuoiodepur S.p.A in San Romano (Pisa, Italy) (43°41’44.5"N, 10°46’02.7"E). *Chaenea vermicularis* sp. nov. (population S4) was retrieved from Marina di Pisa (43□67 □51□□N, 10□27□05□□E) in the province of Pisa (Tuscany, Italy) on the seashore facing the Ligurian Sea. Samples of water and sediment from both Padule di Fucecchio and Marina di Pisa (10-20 mL water up to a total of around 50 mL) were collected in sterile 50-mL Falcon tubes. From Padule di Fucecchio samples were taken at shallow depths (around 20-30 cm from the surface), while samples of sand sediment and algae from Marina di Pisa were recovered at ∼ 1.5 m depth, near a stone breakwater in front of the city embankment. From Cuoiodepur WWTP, bottles of mixed liquors were collected at the outlet of the aeration and anoxic reactors, with *Phialina famelica* sp. nov. retrieved in both the sites. Temperature and salinity were 22.8 °C and 0‰ in Padule di Fucecchio, 23 °C and 33‰ in Marina di Pisa, and 26 °C and 10‰ in Cuoiodepur WWTP. Samples were transferred to the laboratory within 3 h from the collection and three replicate subsamples were obtained from each original sample by stirring and pouring about 10 mL of the liquid into 55 × 14 mm Petri dishes. In the laboratory, each environmental sample was stored at the appropriate salinity level, either in an incubator set at a temperature of 19 ± 1 °C and on a 12:12 h light/dark cycle (light source: NATURAL L36W/76 and FLORA L36W/7 neon tubes, OSRAM, Berlin, Germany) (*Lacrymaria* and *Phialina*), or at room temperature (*Chaenea*). Monoclonal cultures of *Lacrymaria* were established and fed weekly on a mixture of *Paramecium aurelia* and *P. bursaria*. *Paramecium* cultures were in turn fed weekly upon bacteria (*Raoultella planticola* grown in Cerophyll medium) (Boscaro et al., 2013) and the monoclonal culture of the green flagellate *Dunaliella tertiolecta*. *Phialina* and *Chaenea* could not be cultivated in monoclonal cultures and were only maintained for a few (∼ 2 – 3) months thanks to their predating on free-swimming protists present in their own original sample: as for *Phialina*, the presence of *Uronema* sp. both in the samples and in predator’s food vacuoles was sometimes observed in protargol and transmission electron microscopy (TEM) treated cells, while unidentified smaller cells possibly constituted *Chaenea’*s preys according to food vacuoles were sometimes observed in cells processed for TEM.

### Behavioural and morphological analysis

For examination of the swimming behavior, ciliates were observed in a glass depression slide (3 mL) under a dissection microscope (Wild M3, Switzerland) at a magnification of ×12.5 to ×50.0. In order to observe morphological details at a greater magnification and collect photographs, selected cells were placed on glass slides under a differential interference contrast (DIC) microscope with a Leitz (Weitzlar, Germany) instrument equipped with a digital camera (see below) at a magnification of ×300 to ×1250, with the help of a compression device (Skovorodkin, 1990). Feulgen staining with basic fuchsin was used to highlight the nuclear apparatus (Dragesco and Dragesco-Kernéis, 1986; Fokin et al., 2023) of all the three species; the Chatton-Lwoff method of silver nitrate impregnation (Corliss, 1953) and the protargol procedure with silver proteinate (Ji and Wang, 2018) were used to highlight the cortical apparatus and the ciliature of *L. venatrix* sp. nov. and *P. famelica* sp. nov., respectively. Unfortunately, it was not possible to carry out a similar analysis of *C. vermicularis* sp. nov. because the population disappeared from the sample soon after the start of its characterization.

Photomicrographs were captured from appropriate preparations with a digital camera (Canon PowerShot S45), automatically saved as files during optical observation at a magnification of ×500 to ×1250 and used to obtain measurements of living and fixed ciliates. Schematic drawings, based on micrographs of typical living and Feulgen-stained ciliates, were obtained by digitalizing pencil sketches on paper with the vector graphics program Inkscape v.0.92 (https://inkscape.org/); dotted lines were used to represent kineties and inner cell structures.

Terminology and classification based on Lynn (2008) and Vďačný et al. (2011).

### Electron microscopy

For scanning electron microscopy (SEM), ciliates were treated according to Modeo et al. (2006); briefly, selected cells were fixed in 2% osmium tetroxide in 0.1 M cacodylate buffer (pH 7.4), placed on adhesive microscope slides, and dehydrated in an ethanol series. Samples were then attached onto SEM stubs with conductive carbon tape and sputter-coated with gold for SEM observation.

For transmission electron microscopy (TEM), some cells were taken from the cultivation vessels and isolated in fresh medium for some hours. Then, *Lacrymaria* and *Phialina* cells were quickly washed in fresh water and then immediately immersed in a 2.5% (v/v) glutaraldehyde in 0.1 M cacodylate buffer (pH 7.4) 1:1 2% (v/v) osmium tetroxide in distilled water fixing mixture in order to minimize as much as possible cell contraction; *Chaenea* cells were fixed in a 1:1 mixture of 2.5% (v/v) glutaraldehyde in 0.2 M cacodylate buffer (pH 7.4) and 4% osmium tetroxide in distilled water (Fokin et al., 2023). In all cases cells were then processed according to Modeo et al. (2006).

### Whole genome amplification and 18S rDNA sequencing

Starting from a single cell for each of the three species, the total DNA material was amplified via whole-genome amplification (WGA) method, using REPLI-g Single Cell Kit (QIAGEN®, Hilden, Germany). Cells were washed thrice in sterile water at the appropriate salinity and finally in PBS before being transferred in a microfuge Eppendorf for WGA.

On the amplified material Polymerase Chain Reaction (PCR) was performed in a T100™ Thermal Cycler (Bio-Rad, Hercules, CA) to obtain the almost full-length of the 18S rDNA. High-fidelity Takara Ex Taq PCR reagents were employed (Takara Bio Inc., Otsu, Japan) according to the manufacturer’s instructions, using the following primers: 18S F9 (5’-CTG GTT GAT CCT GCC AG-3’) (Medlin et al., 1988) and 18S R1513 Hypo (5’-TGA TCC TTC YGC AGG TTC-3’) (Petroni et al., 2002). PCR cycles were set as follows: 3 min 94 °C, 35×[30 s 94 °C, 30 s 55 °C, 2 min 72 °C], 6 min 72 °C. The so obtained PCR products were purified with the Eurogold Cycle-Pure Kit (EuroClone, Milan, Italy) and subsequently sent for direct sequencing to an external sequencing company (GENEWIZ Germany GmbH (Leipzig, Germany), by adding appropriate internal primers: 18S R536 (5’-CTG GAA TTA CCG CGG CTG-3’), 18S R1052 (5’-AAC TAA GAA CGG CCA TGC A-3’), 18S F783 (5’-GAC GAT CAG ATA CCG TC-3’) (Rosati et al., 2004).

### Illumina sequencing and mitochondrial genome assembly

DNA material belonging to the three organisms, obtained via WGA method was sequenced at GENEWIZ Germany GmbH (Leipzig, Germany), using Illumina NovaSeq (paired-ends 2 × 150 bp) technology to generate 53,558,740 reads for *Lacrymaria* sample, 65,108,368 reads for *Chaenea* sample and 82,362,006 reads for *Phialina* sample.

Preliminary assembly of sequenced reads was performed using SPAdes software (v 3.6.0) (Bankevich et al., 2012). Starting from this preliminary assembly, contigs (i.e. assembled sequences) representing the mitochondrial genome were identified using combining the Blobology pipeline (Kumar et al., 2013), and a tblastn searches using as queries proteins from available mitochondrial genomes belonging to Spirotrichea and Heterotrichea on NCBI. Open Reading Frames (ORFs) on reconstructed mitochondrial genomes were predicted using the GeneMark web server (Besemer and Borodovsky, 2005) setting DNA translation codon table “4” and then manually checked. Ribosomal RNAs and tRNAs were predicted using StructRNAfinder web tool (Arias-Carrasco et al., 2018).

### Phylogenetic analysis

The 18S rDNA sequences of the three organisms were aligned in the ARB software version 5.5 (Westram et al., 2011) on the SSU ref NR102 SILVA database (Quast et al., 2013) using the automatic aligner, followed by manual editing. For the phylogenetic analysis 85 sequences belonging to Litostomatea were selected, plus ten sequences belonging to other Armophorea and Spirotrichea as outgroup. The alignment was trimmed to the shortest sequence obtaining a resulting matrix composed of 1,747 sites. The optimal substitution model was selected with jModelTest 2.1 (Darriba et al., 2012) according to the Akaike Information Criterion. Maximum likelihood (ML) trees were calculated with the PHYML software version 2.4 (Guindon et al., 2010) from the ARB package, performing 1,000 pseudo-replicates. Bayesian inference (BI) trees were inferred with MrBayes 3.2 (Ronquist et al., 2012), using three runs each with one cold and three heated Monte Carlo Markov chains, with a burn-in of 25%, iterating for 1,000,000 generations.

### Symbiont screening

The presence of symbionts associated with the three litostomatean species was verified through 16S rDNA BLAST analysis on the genome assemblies. The three preliminary assemblies obtained from sequenced reads were inspected for the presence of 16S rDNA sequences using Barrnap software V0.5 (Seemann, 2013). The retrieved sequences were checked via blastn analysis on NCBI blast web tool to obtain a preliminary taxonomic annotation. Only 16S rDNAs present on contigs with a coverage higher than 100X were considered as belonging to possible symbionts, while the other ones were classified as contaminants.

Fluorescence *in situ* hybridization (FISH) experiments were performed on *L. venatrix*, using several probe combinations to test the presence of bacteria retrieved from the 16S rDNA screening: the eubacterial generic probes, EUB338I (Amann et al., 1990), and newly designed probes such as Bdel_189 and Bdel_727 (targeting *Bdellovibrio*: 5’-GCTCCTTTGGCCTCAAGAC-3’and 5’-CACGCTTTCGGGAATGAG-3’, respectively), STENO19 (targeting *Alphaproteobacteria*: 5’-TTGCATGTGTTAGGCCTACCG-3’), and VerEup_1174 (targeting *Gammaproteobacteria*: 5’-ACTGACTTGACGTCATCCTCA-3’). Probes were labeled with fluorescein isothiocyanate (FITC) or Cyanine-3 (Cy3) fluorophores. Specimens were fixed in 4% osmium tetroxide (OsO_4_) and, after hybridization procedure (Boscaro et al., 2013), observed with a Nikon ECLIPSE C*i* microscope, equipped with a DS-Ri2 Camera and a D-LEDI Fluorescence LED illumination system (Nikon, Japan). FISH experiments on *P. famelica* and *C. vermicularis* could not be conducted due to the disappearance of the two populations.

## RESULTS

> SYSTEMATICS

> PHYLUM CILIOPHORA DOFLEIN, 1901

> CLASS **LITOSTOMATEA SMALL & LYNN, 1981**

> SUBCLASS **HAPTORIA** Corliss, 1974

> ORDER **LACRYMARIIDA Lipscomb and Riordan, 1990**

> FAMILY **LACRYMARIIDAE DE FROMENTEL, 1876**

> GENUS *LACRYMARIA* BORY DE SAINT VINCENT, 1824

> ***LACRYMARIA VENATRIX* SP. NOV.**

> **(TABLES 1, 2 AND 5; FIGS. 1–3)**

**Table 1.** Morphometrics of *Lacrymaria venatrix* sp. nov.

| Character | Min | Max | Mean | Median | SE | SD | CV (%) | n |
| --- | --- | --- | --- | --- | --- | --- | --- | --- |
| Body length (P) | 73.1 | 121.0 | 92.2 | 88.2 | 4.1 | 16.3 | 17.7 | 16 |
| Body width (P) | 13.5 | 22.5 | 18.9 | 19.7 | 0.6 | 2.4 | 12.9 | 16 |
| Anterior end to Ma, distance (% of body length) (P) | 14.1 | 58.6 | 34.3 | 32.9 | 3.3 | 13.1 | 38.2 | 16 |
| Somatic kineties, number (P) | 17.0 | 18.0 | 17.6 | 18 | 0.2 | 0.5 | 2.9 | 10 |
| Head somatic kineties, number (P) | 10.0 | 15.0 | 13.0 | 13 | 0.5 | 1.5 | 11.6 | 8 |
| Macronucleus length (P) | 17.9 | 35.5 | 27.5 | 27.8 | 1.2 | 4.8 | 17.4 | 16 |
| Macronucleus width (P) | 8.0 | 15.4 | 11.7 | 11.6 | 0.4 | 1.7 | 14.5 | 16 |
| OB length (P) | 2.6 | 5.9 | 4.5 | 4.4 | 0.6 | 1.4 | 30.5 | 5 |
| OB width (P) | 3.6 | 7.1 | 5.8 | 6.1 | 0.6 | 1.3 | 23.2 | 5 |
| Head length (P) | 4.5 | 7.0 | 6.1 | 6.4 | 0.4 | 0.9 | 15.4 | 5 |
| Head width (P) | 6.6 | 8.2 | 7.6 | 7.9 | 0.3 | 0.6 | 8.0 | 5 |
| Oral bulge + head length (P) | 9.3 | 14.3 | 11.4 | 11.5 | 0.3 | 1.2 | 10.8 | 15 |
| Oral bulge + head width (P) | 5.9 | 10.7 | 8.0 | 7.6 | 0.4 | 1.5 | 18.6 | 15 |

Measurements in $\mu\text{m}$ . CV, coefficient of variation; Ma, macronucleus; Max, maximum; Min, minimum; n, number of specimens; OB, oral bulge; P, protargol staining data; SD, standard deviation; SE, standard error. Data on toxicysts length are reported in the text.

### Diagnosis

Body size 73–121 × 14–23 μm after protargol staining with neck retracted, reaching 150-200 μm *in vivo* when contractile neck fully extended. Lanceolate or spindle-shaped trunk, tapering posteriorly, highly contractile. Neck as long as the trunk at full extension. Ovoidal head and dome-shaped oral bulge, both slightly wider than longer. Head and oral bulge can be sometimes completely introverted into the trunk. Macronucleus formed by two ellipsoidal nodules in contact (18–36 × 8–15 μm overall), plus one spherical micronucleus in between. One contractile vacuole, terminal. Ovoidal mucocysts distributed in stripes formed by five to six rows between each pair of adjacent somatic kineties. Toxicysts distributed around the oral region and in the cytoplasm. 10–15 somatic kineties on the head. Dorsal brush separated from the head by a hairless region and consisting of three to six short dikinetids occupying the most anterior region of one to six adjacent somatic kineties. 17–18 somatic kineties covering the trunk, with about 75 monokinetids each. Swimming behaviour: very fast, sometimes sinuous and with retracted neck when they simply move; with extended neck screening the surroundings while hunting; they can exhibit a kind of resting behaviour after eating or after a long period of starving, changing their body shape to a conical or rounded form and remaining immobile, either on the bottom of the container or attached to surfaces. Can form roundish cysts. Found in freshwater.

### Type locality

Padule di Fucecchio (43.807°N, 10.810°E), Fucecchio (Pistoia, Tuscany, Italy); wetland environment (water salinity: 0 ‰).

### Etymology

The specific epithet “*venatrix*” is the feminine singular form of *venator*, Latin for “hunter”, after the voracious predatory behavior of this species.

### Type material

The holotype slide with the silver-stained holotype specimen (indicated with a black circle of ink on the coverslip) and some paratype specimens have been deposited in the collection of the “Museo di Storia Naturale dell’Università di Pisa” (Calci, Pisa, Italy) with registration number CAMUS_2025-1.

### Morphological description

Body length reaching 150-200 μm *in vivo* when contractile neck fully extended (Fig. 1A), while 73–121 × 14–23 μm (mean 92 × 19 μm) after protargol staining with neck retracted (Table 1; Figs. 1A, F); after SEM procedure, about 156 ×13.5 µm in size (Fig. 2A-C). Long, flexible and highly extensible neck arising from the trunk and carrying a head (Figs. 1A, 2A). Neck about as long as the trunk at its full extension, making the whole-body length about 150-200 μm (Fig. 1A). Head less contractile than the neck; trunk shape from lanceolate to almost spherical, with a length-to-width ratio ranging from 4:1 to 5:1, depending on the state of contraction (chemical fixation usually leads to maximal contraction) (Figs. 1A, F; 2B, C).

**Fig. 1.**
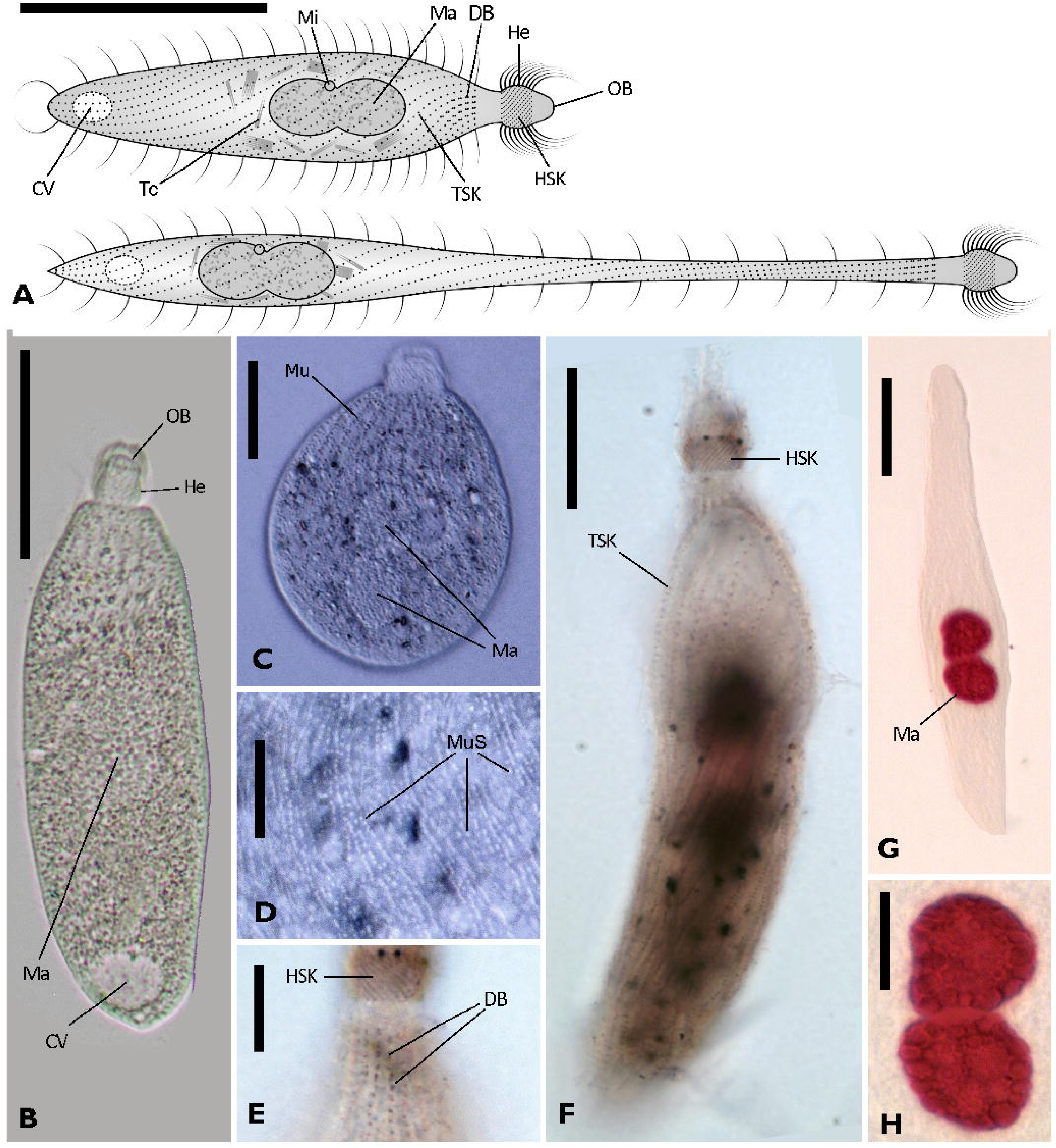
Overall morphology of *Lacrymaria venatrix* sp. nov. A. Schematic reconstruction of a whole cell, with neck contracted (upper draw) and extended (lower draw), based on *in vivo* and fixed cells (Note: toxicysts inserted in the oral bulge are not depicted to avoid overlapping with dorsal brush). B, C. Live pictures showing the whole cell with partially elongated (B) and fully contracted (C) trunk. D. The detail *in vivo* of mucocysts arrangement in stripes. E, F. Protargol-stained specimen, to show details of dorsal brush and head ciliature (E) and somatic ciliature on the trunk (F). G, H. Feulgen-stained specimen with highlighted nuclear apparatus: the whole cell (G) and a closer view (H). CV, contractile vacuole; DB, dorsal brush; He, head; HSK, head somatic kineties; Ma, macronucleus; Mi, micronucleus; Mu, mucocysts; MuS, mucocysts stripes; OB, oral bulge; Tc, toxicysts; TSK, trunk somatic kineties. Scale bars: A, 50 μm; B, C, G, 25 μm; D, H, 10 μm; E, 5 μm; F, 20 μm.

**Fig. 2.**
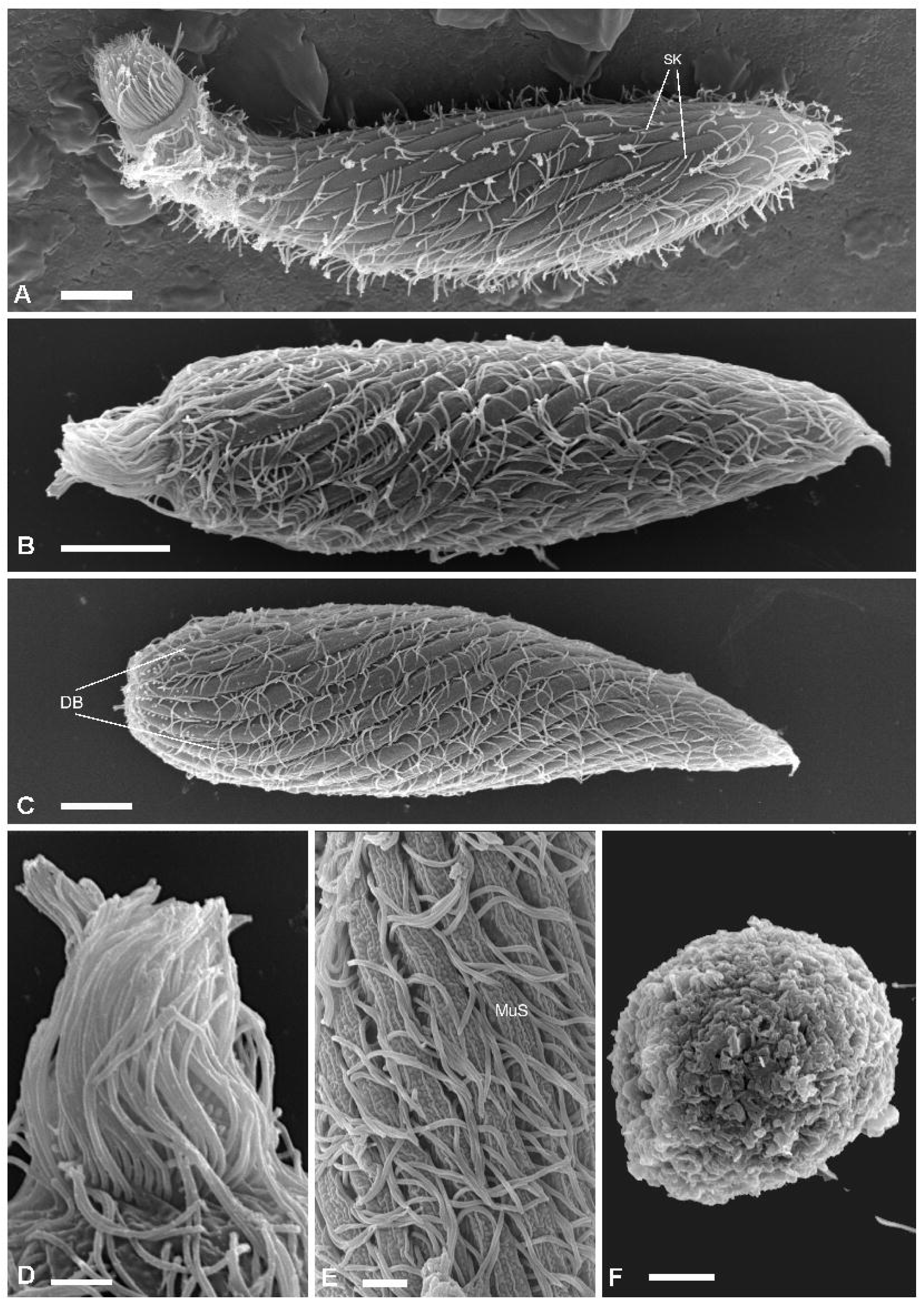
SEM investigation on *Lacrymaria venatrix* sp. nov. A-C. The whole cell, with neck partially extended (A), contracted (B), and with head withdrawn into the trunk (C), respectively (head region always to the left). D. Detail of head and oral bulge, both hidden by head ciliature. E. Detail of the trunk, showing the arrangement of cilia and mucocysts. F. An encysted cell. DB, dorsal brush; MuS, mucocysts stripes. SK, somatic kineties. Scale bars: A-C, 10 μm; D,E, 2 μm; F, 5 μm.

Ovoidal head (4.5–7.0 × 6.6–8.2 μm) visible even at greatest contraction, changing little in shape unlike the trunk (Figs. 1A-C, F; 2A, B, D). Head ends anteriorly in an oral bulge (2.6–5.9 × 3.6–7.1 μm), with a mouth at its apex, surrounded by toxicysts (Fig 3A), which are found distributed in groups consisting in numerous extrusomes in the cytoplasm of the cell central region as well (Figs. 1A, 3C) (see below for ultrastructural details). The oral apparatus is not clearly visible in SEM pictures as it is covered by the ciliature (Fig. 2A-B, D). Head and oral bulge can be sometimes completely introverted into the trunk (Fig. 2C).

**Fig. 3.**
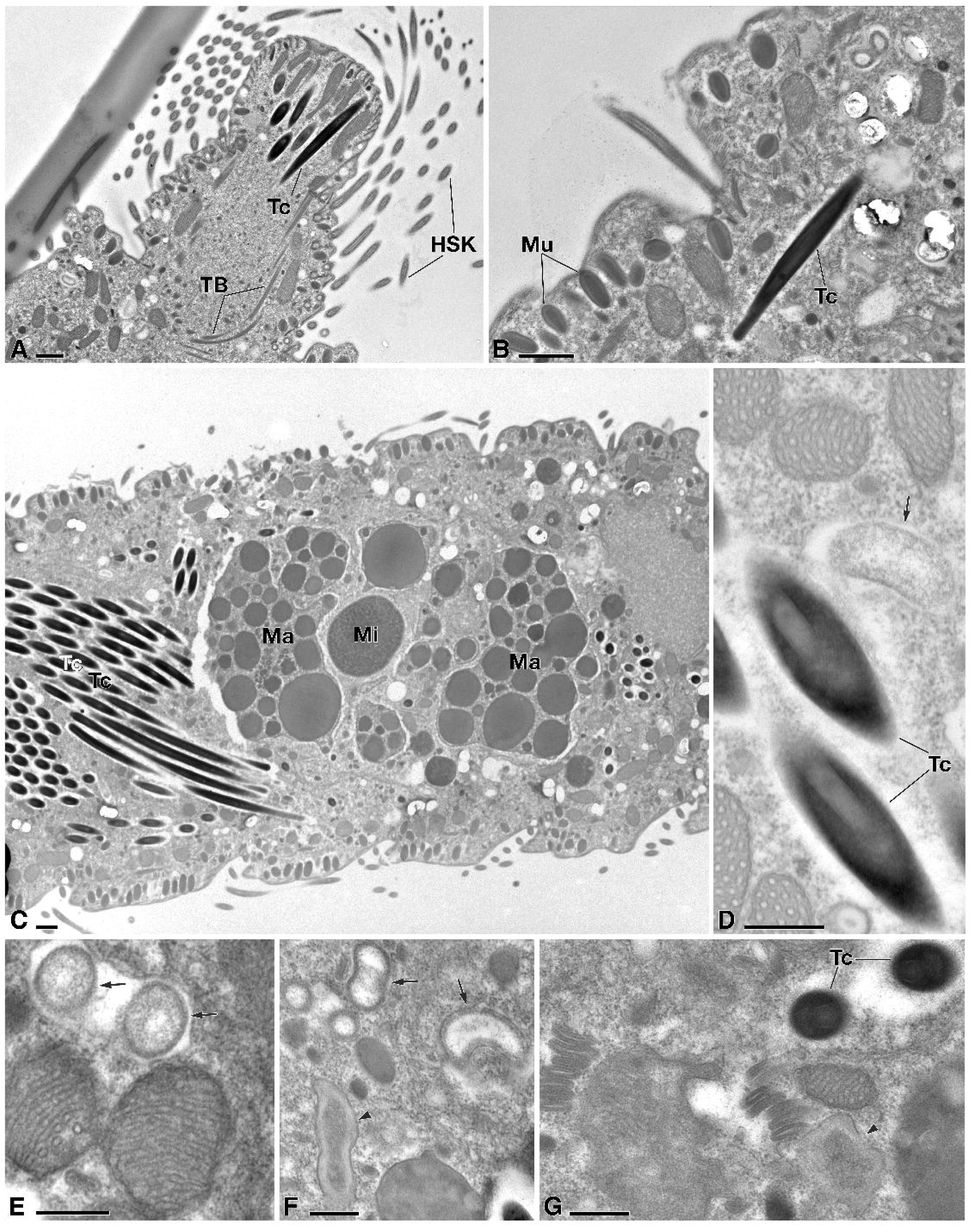
TEM investigation on *Lacrymaria venatrix* sp. nov. A. Detail of head and oral bulge with cilia and toxicysts inserted, and tubulin bundles. B. The cortex with flat alveoli, layered mucocysts and cilia inserted in deep furrows, and some cytoplasm with sparse toxicysts. C. General view of the cytoplasm with several inclusions such as the macronucleus surrounding the roundish micronucleus and flanked by numerous, grouped toxicysts. D-G. Toxicysts and different representatives of the diverse microbiota associated with the ciliate cytoplasm: bean-shaped, electron lucent, double-walled bacteria enclosed in a symbiosome (D-F); irregularly shaped bacteria with and electrondense inner region and a lighter external one (F, G). Arrows indicate bean-shaped bacteria; arrowheads indicate irregularly shaped bacteria; HSK, head somatic kineties; Ma, macronucleus: Mi micronucleus; Mu, mucocysts; TB, tubulin bundles; Tc, toxicysts; Scale bars: A-C, 1 μm; D-G: 0.5 μm.

Macronucleus formed by two spherical nodules in contact (overall size: 18–36 × 8–15 μm), with large vesicles, near centre of the trunk, well-visible after protargol (Fig. 1A, F) and Feulgen staining (Fig. 1 A, G, H), though less obvious (but still visible) *in vivo* (Fig. 1B, C). One spherical micronucleus superimposed over macronucleus nodules (Fig. 1A), not clearly appreciable in Feulgen staining but visible in TEM sections (Fig.3C, see below). One contractile vacuole at posterior end, spherical, about 10 μm in diameter (Fig. 1A, B).

Mucocysts subspherical to ovoidal (see below for details on their ultrastructure), ∼ 0.5–0.7 μm across *in vivo* (Fig. 1D) (0.2–0.3 μm across in SEM treated cells, Fig. 2E); they are densely packed in stripes between ciliary rows (Fig. 2A-C), with five to six rows of them between each pair of kineties (Fig. 2E). This arrangement is visible *in vivo* at high magnification as well (Fig. 1C, D). Each band of mucocysts is ∼ 3.5 μm wide and 2.5 μm apart its neighbours (Fig. 2E). Toxicysts could be observed only in TEM sections (see below).

Head covered by 10–15 dense oblique kineties (Figs. 1A, E, F; 2A, B, D) inserted at ∼ 45° in respect to the antero-posterior axis. Somatic ciliature holotrichous, with 17-18 kineties wrapping in spiral around the trunk (angle depending on the contraction state), set in deep furrows and separated by obvious mucocysts (Figs. 1C-F, 2A-C, E). Somatic kineties on trunk composed of ∼ 75 kinetids, mostly constituted by monokinetids (Figs. 1A, 2A-C). Somatic cilia approximately 8–10 μm long *in vivo*. In the anterior part of the trunk, approximately one to six kineties bear three to six dikinetids, constituting the so-called dorsal brush (Figs. 1A, E; 2C). Dorsal brush separated from the head by a hairless region (Fig. 1A, E, F).

### Notes on the subcellular structure

In the cytoplasm: 1. in the anterior cell region, many tubulin bundles are visible (Fig. 3A), belonging to the chalice-like vortex arrangement of microtubules in the feeding structure and cover functional and structural roles during the feeding, as previously reported (Mikus et al. 2024); 2. there are at least two different types of bacteria, namely an oval to bean shaped one, homogeneous in its inner structure and less electrondense (Fig. 3D-F), and a longer and irregular one, with an inner more electrodense region encircled by a clear region (Fig. 3F, G). Both show a double membrane, but only the first type one is encircled by a clear region and is included in a symbiosome (for further details see the section “Microbial consortium”, below). 3 In the nuclear apparatus the two macronuclear nodules are in contact to each other (∼ 12 µm in diameter) with the single round shaped to ovoid micronucleus (∼ 3.5 µm in diameter) in between, containing a dense chromatin (Fig. 3C). 4. Both mucocysts and toxicysts (Fig. 3A-D, G) appear typical in their general shape and fine structure, in line with literature definitions (Hausmann, 1978; Rosati and Modeo, 2003): mucocysts (∼ 0.6 × 0.3 µm), distributed under the cortex, showing a denser core and a lighter periphery (Fig. 3A-C); toxicysts (up to 12 µm in length, usually 0.4 µm in width) inserted in the apical region and distributed in groups in the cytoplasm; they contain an inner tube encircled by a capsule (Fig. 3A-D, G);

### Occurrence and ecology

During the permanence in the laboratory, *L. venatrix* strain PDF5b_Lac3 grew by predating on *Paramecium aurelia* and *Paramecium bursaria*. In the laboratory, the ciliate displayed at least three distinct motor behaviours: 1. fast swimming with a retracted neck, characterized by straight and rapid movement; 2. hunting behaviour, involving slow swimming with frequent back-and-forth movements, always with the neck extended and actively scanning the surroundings in all directions; 3. resting state, observed after feeding or following several days of starvation, during which cells appeared roundish or conical, remained immobile, and were attached to the bottom or to various surfaces. This resting state could be interrupted by a simple mechanical stimulus, which caused the cell to abruptly change shape and resume swimming. The production of roundish cysts (∼ 17 µm in diameter) was observed (Fig. 2F) after three weeks of starvation.

### Gene sequence

The 18S rDNA sequence of *L. venatrix* (strain PDF5b_Lac3) obtained from PCR resulted 1,608 bp long, and it has been deposited in NCBI GenBank database with the accession number PQ628060. The 18S rDNA sequence of *L. venatrix* showed the highest identity value with the sequence of *Phialina vertens* (MF474348): 98.38% (2 gaps, 24 mismatches).

### Mitochondrial genome

The mitochondrial genome of *L. venatrix* resulted in a single linear molecule of 62,315 bp (for further details see the “Mitochondrial genome” section, below).

### Microbial consortium

The screening of the preliminary assembly for bacterial 16S rDNA sequences revealed four different bacteria (OTUs #1–4) putatively associated with *L. venatrix.* We found four different 16S rDNA sequences, belonging to different bacterial classes and phyla: one *Bdellovibrio* sp. (*Bdellovibrionota, Bdellovibrionia*), OTU #1, two *Gammaproteobacteria* (*Pseudomonadota*), OTU #2 and #4; and one *Alphaproteobacteria* (*Pseudomonadota*), OTU #3. FISH results were negative for all the probes tested (data not shown). Details about sequences length, coverage, accession numbers, and best BLAST hits are given in Table 2. For ultrastructural details, see above.

> SUBCLASS **HAPTORIA** Corliss, 1974

> ORDER **LACRYMARIIDA Lipscomb and Riordan, 1990**

> FAMILY **LACRYMARIIDAE DE FROMENTEL, 1876**

> GENUS *PHIALINA* BORY DE SAINT VINCENT, 1824

> ***PHIALINA FAMELICA* SP. NOV.**

> **(TABLES 2, 3 AND 6; FIGS. 4–6)**

**Table 2.**
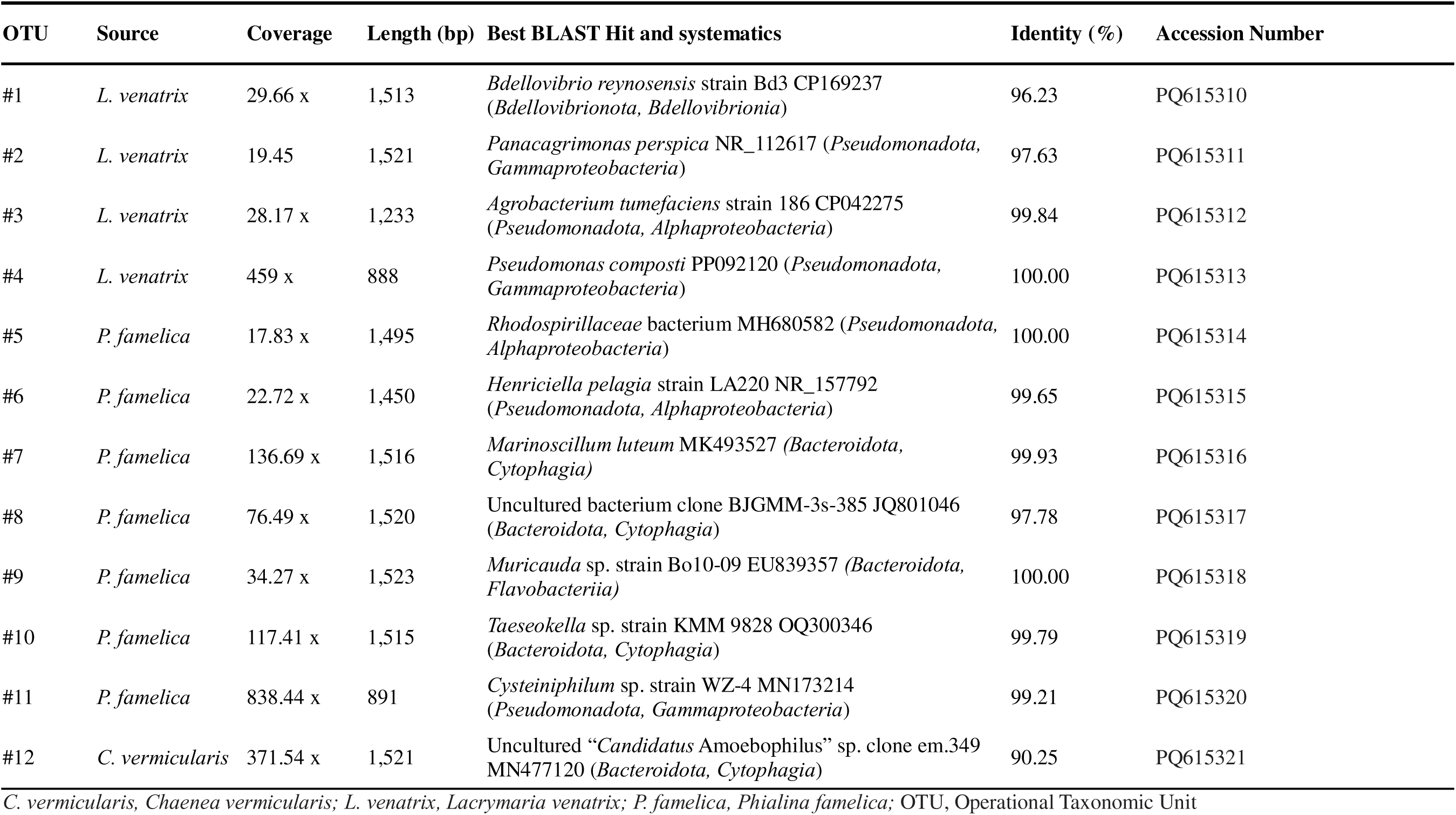
16S rDNA sequences of prokaryotes associated to the three studied litostomatean ciliates.

### Diagnosis

Body size *in vivo* 73–99 × 20–33 μm, cylindrical-to-lanceolate trunk, rounded posteriorly, moderately contractile; no distinct neck or neck-like region, as the head is in continuity with the trunk. Barrel-shaped head and dome-shaped oral bulge, each about as wide as long. Single macronucleus, elongated ellipsoidal or reniform (15–40 × 3–7 μm, after protargol staining), one elliptical micronucleus adjacent to macronucleus. One contractile vacuole, terminal. Spherical to elliptical mucocysts distributed in stripes arranged in about five parallel rows between pair of adjacent somatic kineties. Toxicysts of three different morphological types inserted around the oral region; toxicysts of the two longer types scattered in the cytoplasm of the trunk. About 8-9 kineties on the head. Dorsal brush in continuity with both head and somatic kineties and consisting of four rows including four to five dikinetids each. 12–16 somatic kineties on trunk, with about 55 monokinetids each. Movement by fast swimming along a helical trajectory while rotating around the main body axis. Found in brackish water, both in oxic and anoxic conditions.

### Type locality

Cuoiodepur Wastewater Treatment Plant (43°41’44.5"N, 10°46’02.7"E), San Miniato, (Pisa, Tuscany, Italy) (water salinity 5–10 ‰)

### Etymology

The specific epithet “*famelica*” is the feminine singular form of *famelicus*, Latin for “hungry” or “voracious”, after the active predatory behavior of this species.

### Type material

The holotype slide with the silver-stained holotype specimen (indicated with a black circle of ink on the coverslip) and some paratype specimens have been deposited in the collection of the “Museo di Storia Naturale dell’Università di Pisa” (Calci, Pisa, Italy) with registration number CAMUS_2025-2.

### Morphological description

Body length *in vivo* 73–99 × 20–33 μm (mean: 88 × 25 μm), composed of a slightly contractile head and a moderately contractile trunk (Table 3); trunk shape is cylindrical with blunt or oblanceolate posterior end, with a length-to-width ratio ranging from 3:1 to 5:1 (mean 3.7:1), depending on the state of contraction (Figs. 4A-E, 5A). Chemical fixation leads to a contraction of 30–50 % in length: protargol-stained specimens reduced to 35–66 × 15–34 μm, (mean: 48 × 23 μm), with a length-to-width-ratio 2.2:1 and a shape elliptical to ovate compared to *in vivo* individuals (Fig. 4A-D); SEM processed cells ∼ 46 ×12 µm (Fig. 5A), with a cell shrinkage of 58% and 50% compared to *in vivo* cell length and width, respectively.

**Fig. 4.**
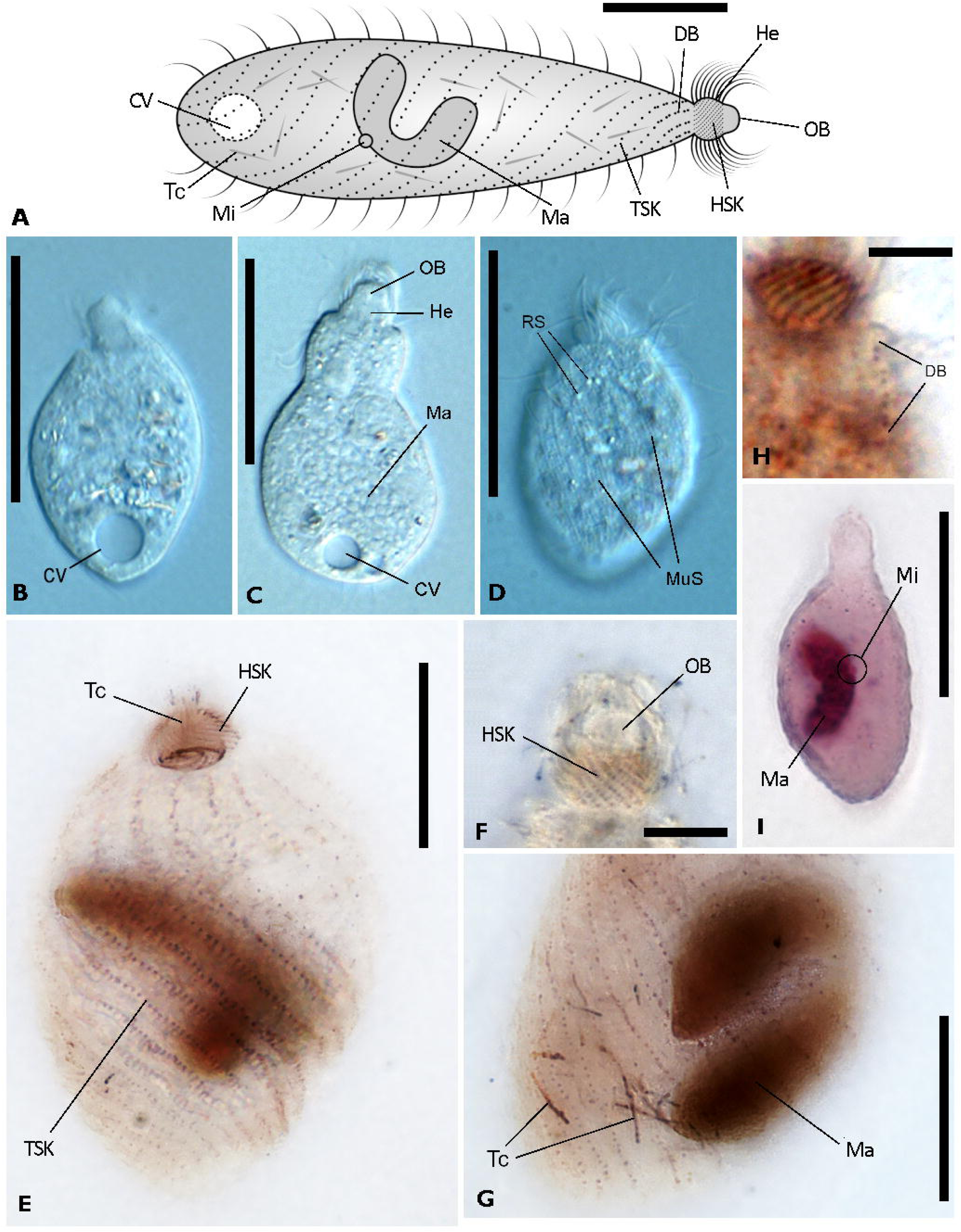
Overall morphology of *Phialina famelica* sp. nov. A. Schematic reconstruction of a whole cell based on *in vivo* and fixed cells (Note: toxicysts inserted in the oral bulge are not depicted to avoid overlapping with dorsal brush). B, C, D. *In vivo* observation of whole cells at different contraction states (B, C); refractive structures and mucocysts distributed in stripes between the kinetids are shown (D). E-H. Protargol-staining: a whole cell (E) with toxicysts inserted in the oral bulge, details of head ciliature (F), cell posterior with cytoplasmic toxicysts (G), and dorsal brush (H). I. Feulgen-stained whole specimen with macronucleus and micronucleus highlighted. CV, contractile vacuole; DB, dorsal brush; He, head; HSK, head somatic kineties; Ma, macronucleus; Mi, micronucleus; MuS, mucocysts stripes; OB, oral bulge; RS, refractive structures; Tc, toxicysts; TSK, trunk somatic kineties. Scale bars: A, 20 μm; B-D, 30 μm; E, G, I, 20 μm; F, H, 5 μm.

**Fig. 5.**
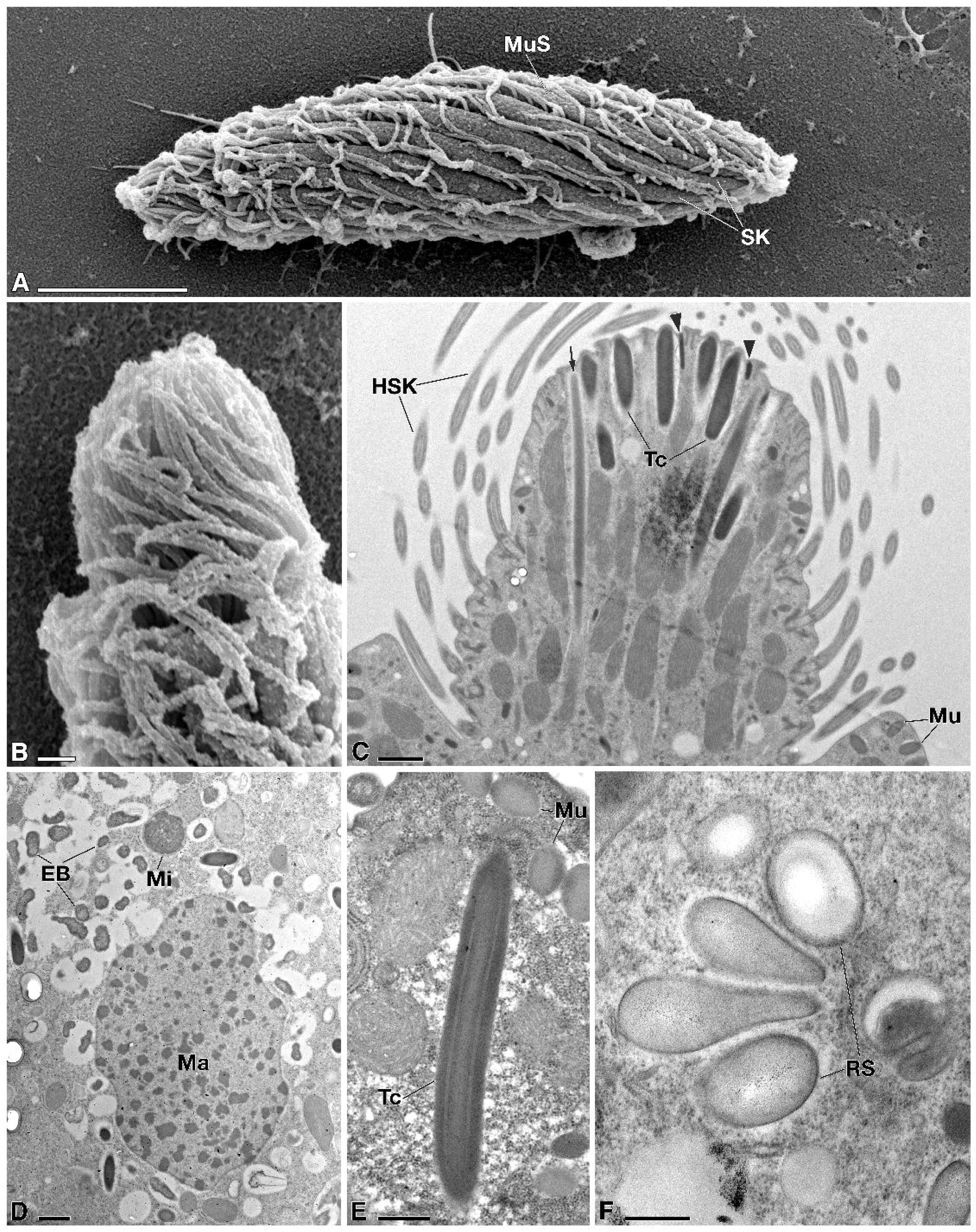
SEM and TEM investigation on *Phialina famelica* sp. nov. A, B. SEM investigation: a whole cell, with head region to the left (A), and detail of head and oral bulge, both hidden by head ciliature (B). C-F. TEM investigation: detail of head and oral bulge with cilia, three different types of toxicysts (see text for explanation), mucocysts, and mitochondria (C); a general view of the cytoplasm with several inclusions such as micronucleus and macronucleus, surrounded by numerous, homogeneously electrondense, irregularly shaped bacteria, encircled by a white halo (D); detail of the inner layered structure of toxicysts type II and mucocysts (E); detail of the refractive structures (F). Arrow indicates toxicyst type I; arrowheads indicate toxicyst type III; EB, electrondense bacteria; HSK, head somatic kineties; Ma, macronucleus; Mi, micronucleus; Mu, mucocysts; MuS, mucocysts stripe; RS, refractive structures; Tc, toxicysts type II; SK, somatic kineties. Scale bars: A, 10 μm; B-D, 1 μm; E, F, 0.4 μm.

**Table 3.**
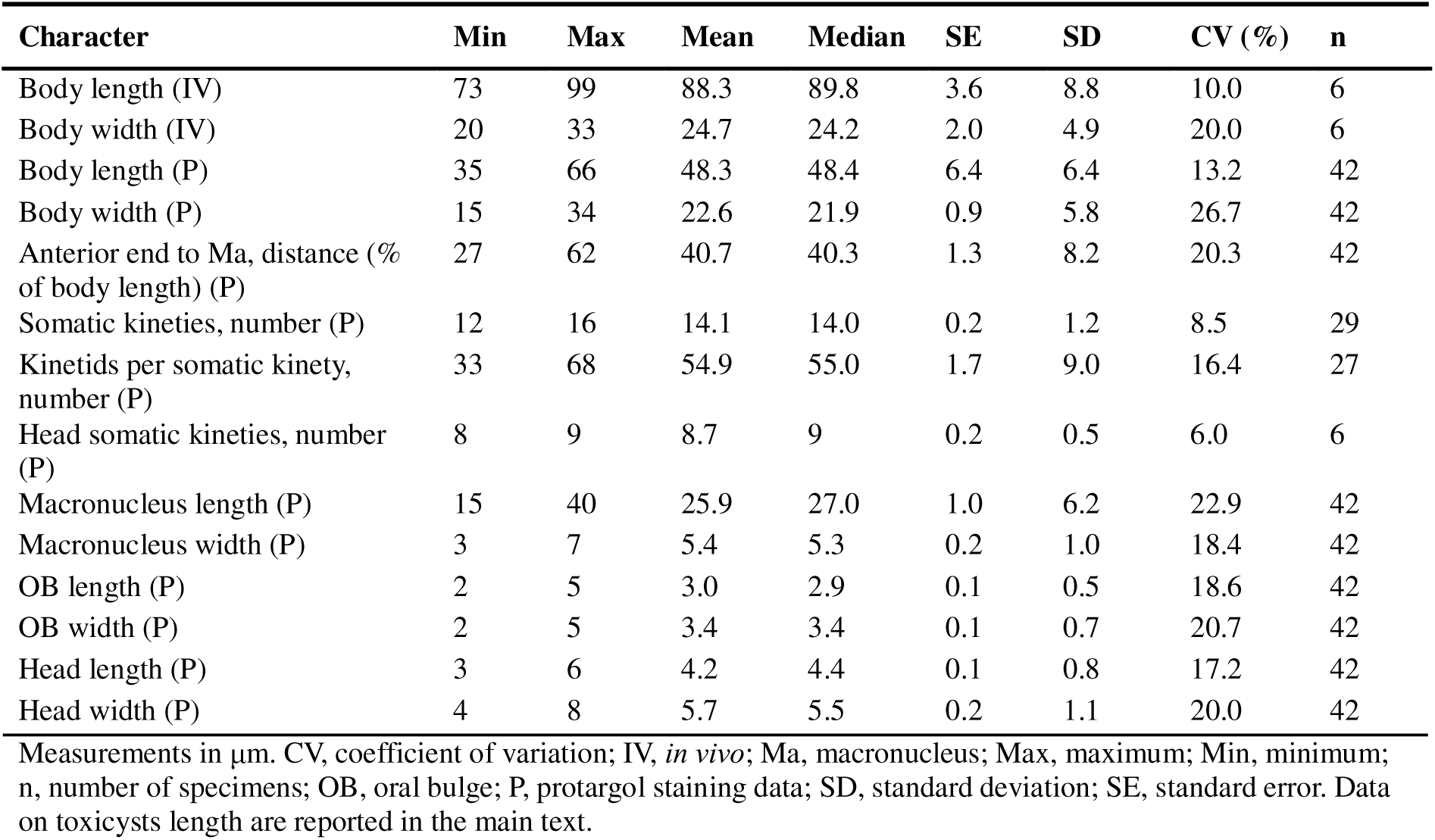
Morphometrics of *Phialina famelica* sp. nov.

Barrel-shaped head (6.4 × 6.2 μm *in vivo*) directly attached to trunk (i.e., no neck is present in between), with a hemispherical oral bulge (2.5 × 4.6 μm) at its anterior end (Figs. 4A-F, 5A-C) bearing a mouth at its apex. Head sometimes slightly retracted into trunk under high compression or chemical stimulation, generally hidden by a dense ciliature in SEM view (Fig. 5A, B). Oral apparatus occupying about 15% of body length in protargol-stained specimens, posteriorly delimited by circumoral kinety (Fig. 4E), with tips of toxicysts inserted in the oral bulge as typical of the genus (Figs. 4E, 5C).

One macronucleus reniform (Fig. 4A) or elliptical (Fig. 4C) *in vivo* (17–38 × 7–15 μm, mean: 27 × 10 μm, width-to-length ratio ∼ 1:3), often appearing oblong and curved after fixation and staining (Fig. 4A, E, G, I). After protargol staining, macronucleus 15–40 × 3–7 μm (mean: 26 × 5 μm) in size (Fig. 4E). Micronucleus hard to recognize both *in vivo* and in stained specimens; single elliptical micronucleus, ∼ 1 μm in length, adjacent to macronucleus, evident in Feulgen-stained specimens (Fig. 4I) and in TEM processed cells (see below; Fig. 5D). One contractile vacuole at posterior end (Fig. 4A-C).

In protargol-stained cells, rod-shaped toxicysts (∼8.5 μm in length) densely attached to the oral region (Fig. 4E) appear weakly responsive to the staining compared to those scattered in the trunk, which are more highlighted (Fig. 4A, G); but, apparently, more than one type of toxicysts, different in size and thickness, are present according to TEM observation (Fig. 5C; see below). Mucocysts spherical to elliptical, ∼ 0.2–0.4 μm across oriented perpendicularly to cell surface, densely and regularly arranged in stripes of ∼ 5 parallel rows between adjacent ciliary rows (Figs. 4D, 5A), less distinct than in *Lacrymaria*. Cytoplasm colourless, usually packed with regularly shaped refractive inclusions (Fig. 4D). The latter, together with mucocysts, are more appreciable in TEM-processed specimens (see below; Fig. 5E, F).

Head covered by 8–9 densely arranged kineties running spirally (Figs. 4A, 5A, B); each kinety composed of densely arranged monokinetids, poorly visible in protargol staining (Fig. 4E, F). Somatic ciliature holotrichous, with 12–16 kineties, helically or parallelly inserted respect to body axis depending on the state of cell contraction (Figs. 4A, E, 5A). Somatic kineties composed of ∼ 55 monokinetids (Figs. 4A, E, 5A); in four of them, three to four dikinetids in their anterior part form the dorsal brush, inconspicuous both *in vivo* and after protargol staining (Fig. 4A, H) and SEM, posteriorly to which the monokinetids extend. Somatic cilia ∼ 7 μm long *in vivo*.

### Notes on the subcellular structure

In the cytoplasm, numerous inclusions were observed (Fig. 6) such as diffused oval to elongated refractive inclusions (∼ 1.5 μm in length) (Fig. 5F), lipid droplets (∼2 μm in diameter) (Fig. 6D), and food vacuoles of different sizes (Fig. 6B) with putative residuals of prey. Additionally, we could observe at least four types of bacteria with different morphology, which are representatives of the variegated ciliate-associated microbiome: 1. homogeneously electrondense, irregularly shaped bacteria, encircled by a white halo (Figs. 5D, 6A, C), which were especially abundant around the macronucleus; 2. flagellated bacteria encircled by a white halo (Fig. 6A, C), which sometimes showed virus-like particles in their cytoplasm; 3. almond-shaped bacteria, with a lighter cytoplasm and a thinner white halo around (Fig. 6B); 4. roundish bacteria, rich in ribosomes, enclosed in symbiosomes tightly associated with rough endoplasmic reticulum cisternae (Fig. 6D). Finally, in *P. famelica* we could observe two different types of extrusomes: toxicysts and mucocysts (diam: ∼ 0.4 µm) (Figs. 5C, E, 6A, B, D). However, according to TEM investigation, there likely be three different types of toxicysts which alternate each other while inserting in the oral bulge (Fig. 5C): 1. a slightly electrondense, long and slender type (type I) (size: ∼ 6.5 × 0.2 µm); 2. a middle-sized type (type II) (size: ∼ 3.5 × 0.4 µm); and 3. a very short and electrondense type (type III) (size: ∼ 0.7 × 0.1 µm). Type II toxicysts are also visible scattered in the cytoplasm in TEM sections (Figs. 5D, E, 6D); together with the mucocysts (∼ 0.4 × 0.3 µm), they appear very typical in their general shape and fine structure, according to literature descriptions of the two kinds of extrusive organelles (Hausmann, 1978; Rosati and Modeo, 2003): the toxicysts include a tube and a capsule (Figs. 5C, E, 6D); the mucocysts show a denser core and a lighter periphery (Figs. 5C, E, 6A, B). It was not possible to precisely disclose which of the two longer toxicysts types (i.e., type I or type II) corresponds to the extrusomes observed under the light microscope after protargol staining (Fig.4 E, G), which apparently does not adequately stain all the toxicysts types; nevertheless, according to the size, we hypothesized it should be type I.

**Fig. 6.**
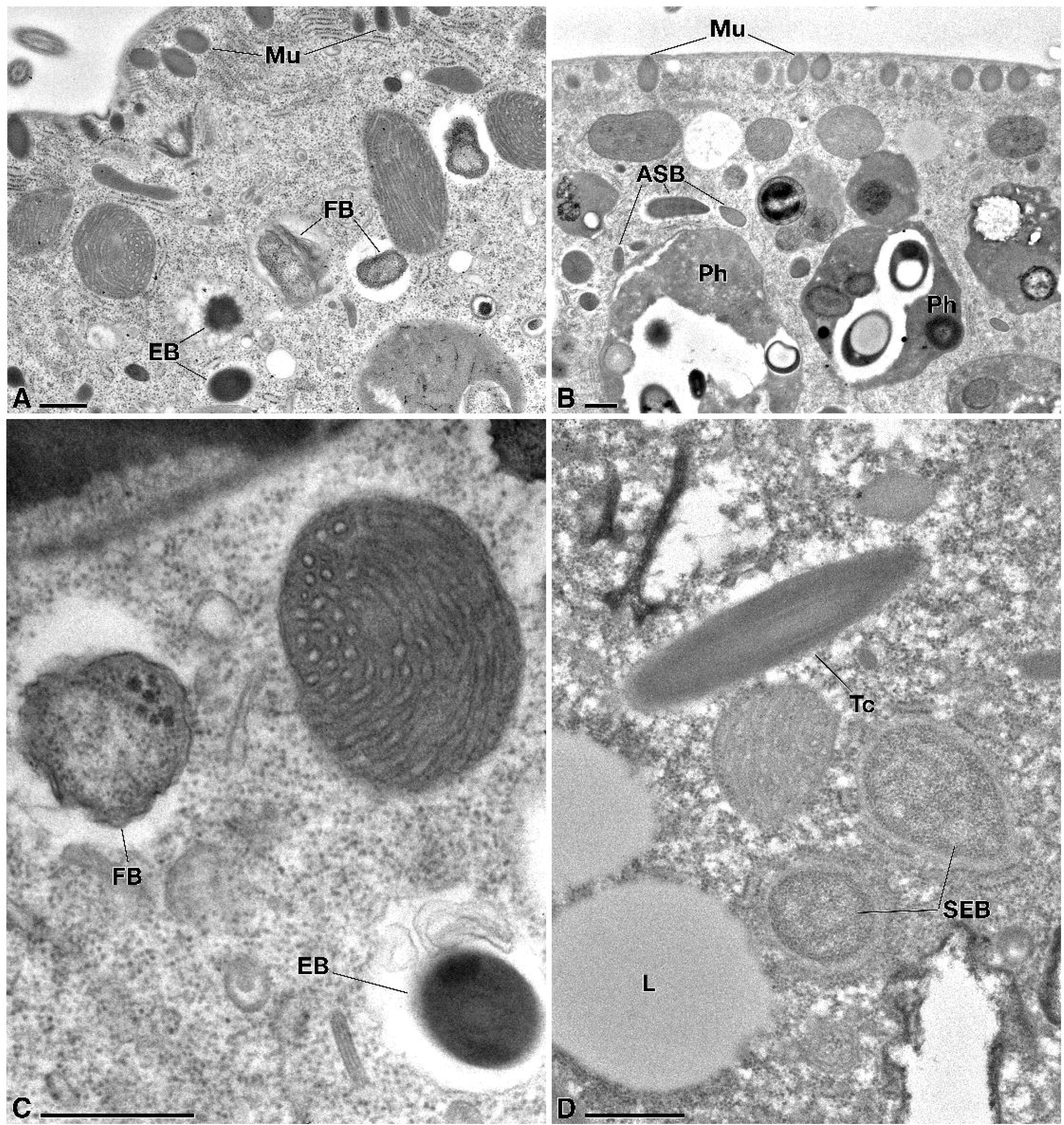
TEM investigation on *Phialina famelica* sp. nov. Focus on the ciliate cytoplasm content, including different representatives of the ciliate-associated diverse microbiota. A. Flagellated bacteria encircled by a white halo and electrondense roundish bacteria. B. Almond-shaped bacteria with a low electrondense cytoplasm and a thinner white halo around. C. Closer view of the flagellated bacteria, which, sometimes, show virus-like particles in their cytoplasm. D. Two roundish bacteria, rich in ribosomes, enclosed in symbiosomes tightly associated with rough endoplasmic reticulum cisternae. ASB, almond-shaped bacteria; EB, electrondense bacteria; FB, flagellated bacteria; Ph, phagosome; L, lipid reserve drop; Mu, mucocysts; SEB, symbiosome-enclosed bacteria; Tc, toxicysts type II. Scale bars: A-D: 0.5 μm.

### Occurrence and ecology

During the permanence in the laboratory, the *P. famelica* population grew by predating on other ciliates such as *Uronema* sp. In the laboratory, cells swam fast along a helical trajectory by rotating around the main body axis. No cyst formation was observed.

### Gene sequence

The 18S rDNA sequence of *P. famelica* obtained from PCR resulted 1,605 bp long, and it has been deposited in NCBI GenBank database with the accession number PQ628061. The 18S rDNA sequence of *P. famelica* showed the highest identity with the sequence of *Phialina salinarum* (EU242508): 97.58% (6 gaps, 33 mismatches).

### Mitochondrial genome

The mitochondrial genome of *P. famelica* resulted in a single linear molecule of 52,439 bp (for further details see the “Mitochondrial genome” section, below).

### Microbial consortium

The screening of the preliminary assembly for bacterial 16S rDNA sequences revealed seven different bacteria (OTUs #5–11) putatively associated with *P. famelica*. We found seven different 16S rDNA sequences, belonging to different bacterial classes and phyla: two *Alphaproteobacteria* and one *Gammaproteobacteria* (*Pseudomonadota*), OTUs #5-#6 and #11; three *Cytophagia* and one *Flavobacteriia* (*Bacteroidota*), OTUs #7-10. Details about sequences’ length, coverage, accession numbers, and best BLAST hits are given in Table 2. For the ultrastructural details, see above.

> SUBCLASS **HAPTORIA** Corliss, 1974

> *INCERTAE SEDIS*

> FAMILY **CHAENEIDAE KWON ET AL., 2014**

> GENUS *CHAENEA* QUENNERSTEDT, 1867

> *CHAENEA VERMICULARIS* SP. NOV.

> **(TABLES 4 AND 7; FIGS. 7–9)**

**Table 4.** Morphometrics of *Chaenea vermicularis* sp. nov.

| Character | Min | Max | Mean | Median | SE | SD | CV (%) | n |
| --- | --- | --- | --- | --- | --- | --- | --- | --- |
| Body length (F) | 98.0 | 450.0 | 192.1 | 189.3 | 29.4 | 106.0 | 55.2 | 13 |
| Body width (F) | 10.5 | 15.6 | 13.8 | 13.1 | 0.4 | 1.4 | 10.1 | 13 |
| Body length (SEM) | 93.0 | 140.0 | 117.2 | 122.2 | 9.4 | 20.6 | 22.0 | 5 |
| Body width (SEM) | 7.0 | 15.0 | 12.3 | 13.30 | 1.3 | 3.0 | 25.0 | 5 |
| Somatic kineties, number (IV) | 8.0 | 12.0 | 10.5 | 10.0 | 0.2 | 1.0 | 10.0 | 20 |
| Somatic cilia length (IV) | 4.0 | 5.3 | 4.4 | 4.1 | 0.1 | 0.3 | 6.6 | 20 |
| Rows of mucocysts per strip, number (IV) | 5.0 | 7.0 | 5.5 | 5.2 | 0.2 | 1.0 | 18.9 | 18 |
| Toxicysts length (IV) | 6.6 | 7.0 | 6.9 | 6.5 | 0.1 | 0.6 | 8.7 | 15 |
| Anterior body end to DB, distance (IV) | 10.5 | 13.0 | 11.7 | 11.3 | 3.1 | 8.1 | 69.2 | 7 |

Measurements in $\mu\text{m}$ . 1 Measurements were taken in the equatorial part of the cell body. CV, coefficient of variation (%); DB, dorsal brush; F, Feulgen staining; IV, *in vivo*; Ma, macronucleus; Max, maximum value; Mean, mean value; Min, minimum value; n, number of specimens; SEM, scanning electron microscope; SD, standard deviation; SE, standard error.

### Diagnosis

Body size *in vivo* 130–520 × 11–17 µm, elongated to vermiform, cylindrical anteriorly and slightly tapering to rounded posteriorly, only slightly contractile. Cell anterior end sometimes somewhat bent with a slightly protruding oral bulge on its apex. Macronucleus consisting of a huge number of nodules which, depending on the phase in the ciliate life cycle, can include either only one type nodules, i.e., spherical to ovoidal in shape (∼ 3–4 µm in diameter), observed after division process, or two types, i.e., smaller irregular (1–2 µm-sized but variable; numerous) and relatively larger rounded ones (∼ 5 µm-sized, always rarer), observed before division process; possibly several micronuclei, difficult to observe due to the overlapping with macronuclear nodules. One contractile vacuole without collecting channels in the posterior cell end. Roundish to oval mucocysts distributed in stripes and arranged in 5-7 parallel rows between pair of adjacent somatic kineties. Numerous mace-like toxicysts inserted in the oral bulge and sparse in the cytoplasm. Dorsal brush consisting of 4 rows of dikinetids, smaller than the somatic cilia and distributed in a zigzag pattern, in continuity with the head somatic kineties towards the anterior cell end, and with 8–10 somatic kineties composed of monokinetids towards the posterior cell end.

### Type locality

Coastline of the Ligurian Sea (43°40′20″N 10°16′37″E), Marina di Pisa (Pisa, Tuscany, Italy) (water salinity: 33 ‰).

### Etymology

The specific epithet “*vermicularis*” is the adjectival form of *vermiculus*, diminutive of *vermis*, Latin for “worm”, with reference to the long and slim shape of the cell.

### Type material

One stub of SEM treated specimens from the type population (S4) of *Chaenea vermicularis* has been deposited in the collection of the Unit of Zoology-Anthropology of the Department of Biology at Pisa University (Pisa, Italy) with registration number UNIPI_2025_1.

### Morphological description

Cell shape elongated to vermiform; body cylindrical anteriorly and slightly tapering to rounded posteriorly (Figs. 7, 8A). Body size *in vivo*: ∼130–520 × 11–17 µm (Fig. 7A, D) and after Feulgen staining, 98–450 × 10.5–15.6 µm (mean ∼192 μm × 14 μm) (Fig. 8A) (Table 4). After SEM procedure, 93–140 × 7–15 µm (mean 117.2 μm × 12.3 μm) (Fig. 7F, G) with a cell shrinkage of 52% and 11% compared to *in vivo* length and width, respectively (Table 4). Body length-to-width ratio different between mature (i.e., proximate to division) and young (i.e., freshly divided) individuals: in the former, ratio 23:1; in the latter, ratio close to 10:1. Cell body only slightly contractile. Cell anterior end sometimes slightly bent (Figs. 7D, E, G, 8A). Oral bulge slightly protruding on the cell anterior end, with the mouth at its apex (Fig. 7C). Rounded oral bulge occupying the anterior body end, ∼ 3–4 µm in diameter *in vivo* (Fig. 7A-E), not visible in SEM-treated cells due to the covering by oralized somatic and circumoral cilia (Fig. 7F-H) with which it is in continuity.

**Fig. 7.**
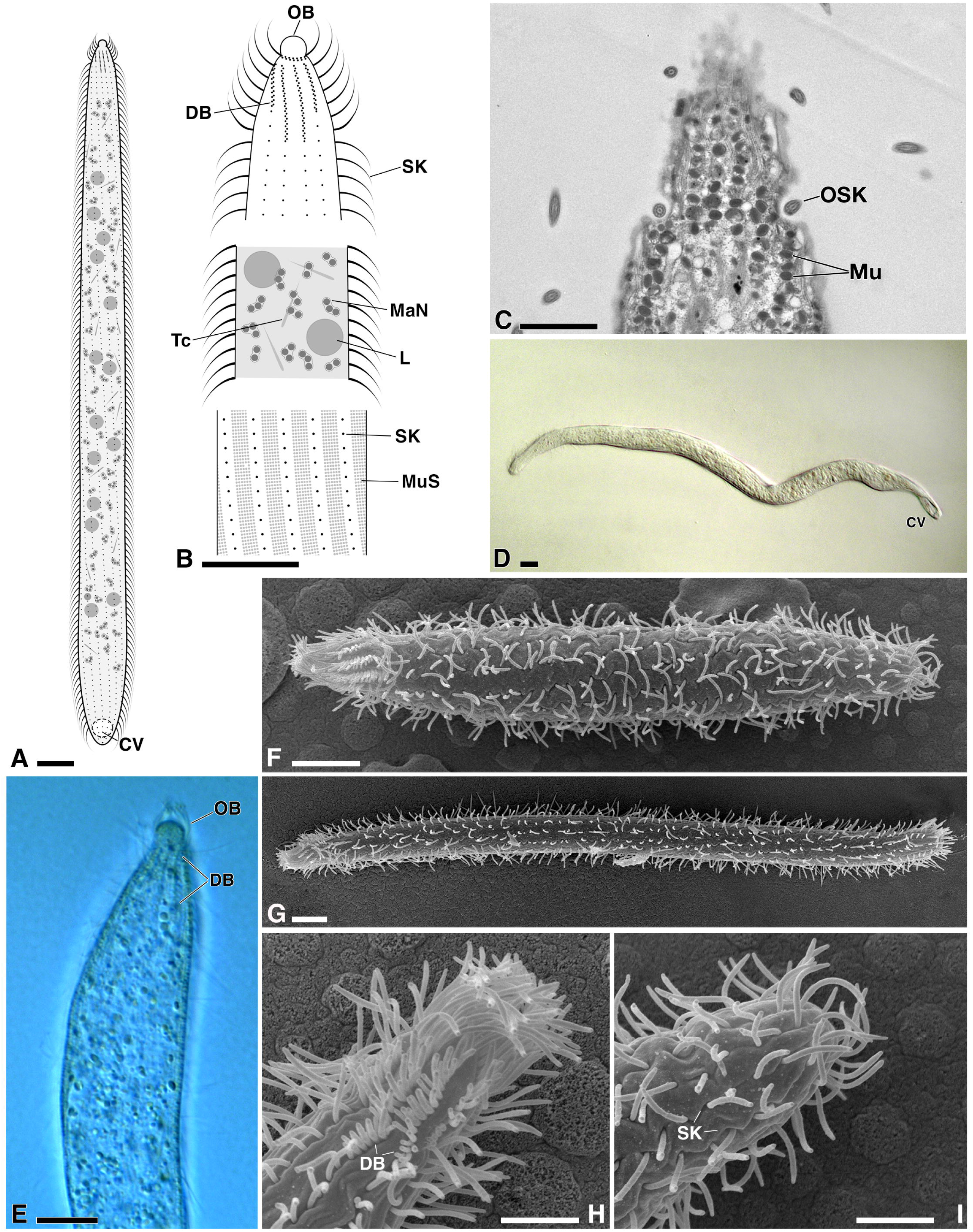
Overall morphology of *Chaenea vermicularis* sp. nov.. A, B. Schematic reconstruction based on *in vivo* and fixed cells of a whole cell (A) and magnifications to show cell details, i.e., the cell anterior and the dorsal brush, some typical cytoplasm content, and the surface structure with the somatic ciliary rows and the mucocysts stripes of the trunk (B) (Note: toxicysts inserted in the oral bulge are not depicted to avoid overlapping with dorsal brush). C, TEM investigation: cell anterior region sectioned at a superficial level to show its silhouette. D-E. *In vivo* investigation: a ciliate in its full length, with the terminal contractile vacuole (D); detail of the anterior cell portion (E). F-I. SEM investigation: general shape of a small (F) and a middle-sized (G) specimens, respectively (cell anterior region always to the left); a closer view of the cell anterior from the dorsal side, with the four rows of shorter cilia arranged in a zig-zag pattern, forming the dorsal brush (the two central rows are pointed out); note that the rounded oral bulge is not visible due to been covered by somatic and circumoral cilia placed anteriorly to the dorsal brush (H); a closer view of the posterior cell end (I). CV, contractile vacuole; DB, dorsal brush; L, lipid reserve drop; MaN, macronuclear nodule; Mu, mucocysts; MuS, mucocysts stripes; OB, oral bulge; OSK, oralized somatic kineties; SK, somatic kineties; Tc, toxicyst. Scale bars: A-B, D-I,10 µm; C, 2 µm.

**Fig. 8.**
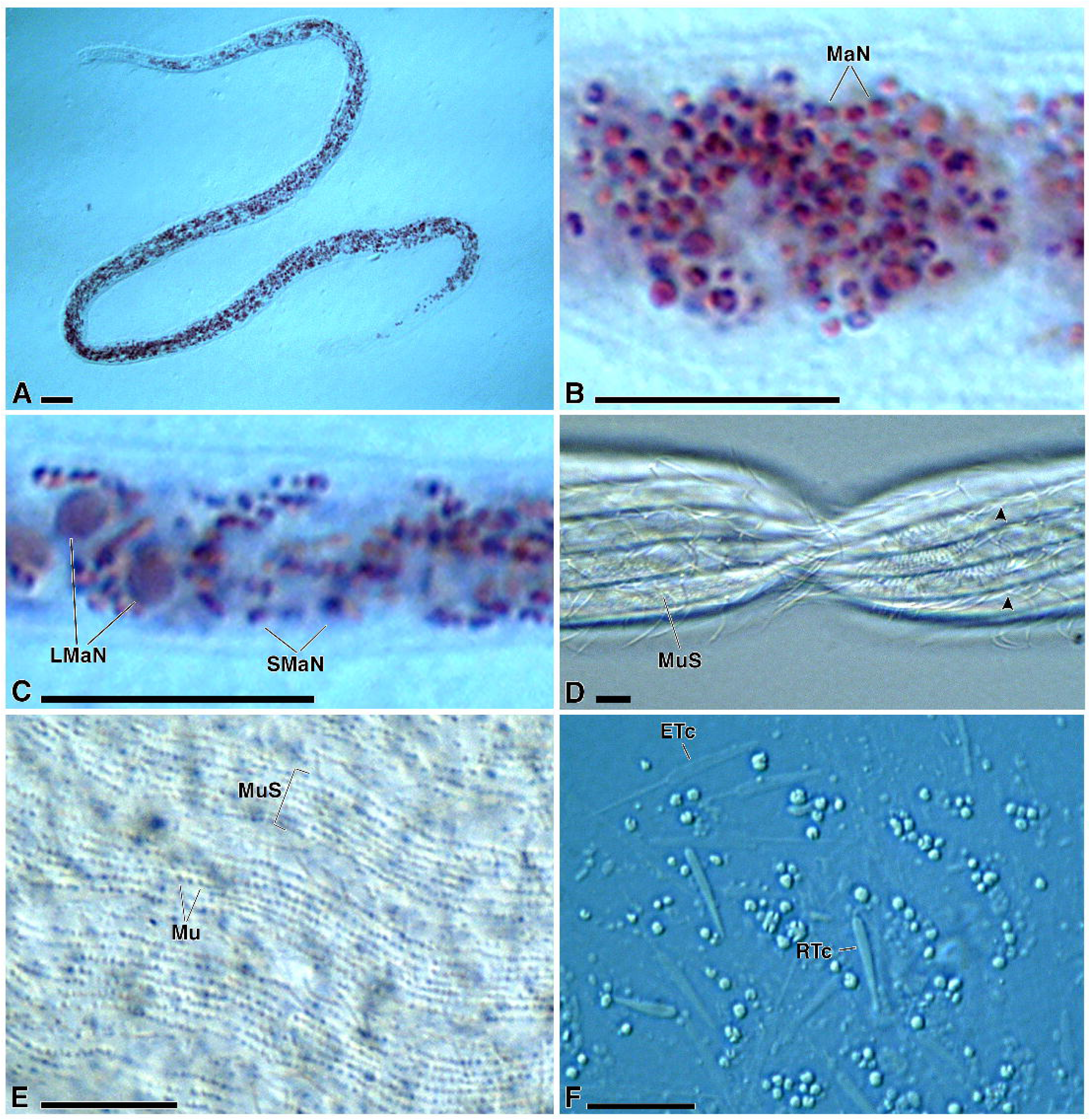
Morphological details of fixed and living *Chaenea vermicularis* sp. nov. cells. A-C. Feulgen staining, highlighting numerous macronuclear nodules distributed in the cytoplasm: general view of a cell (A); the cytoplasm of a specimen after cell division with macronuclear nodules homogeneous in shape and sizes (B); the cytoplasm of a specimen before cell division (C) with two different subtypes (i.e., larger and smaller) of macronuclear nodules. D, E. View of living cells: the somatic kineties of a dividing specimen (D) alternate with the stripes formed by the mucocysts (D, E). F. *In vivo* observation of the cytoplasm of a squashed cell: intact (i.e., resting state) and extruded toxicysts are present. Arrowheads indicate somatic kineties; ETc, extruded toxycist. LMaN, larger macronuclear nodules; MaN, macronuclear nodules; Mu, mucocysts; MuS, mucocysts stripe; RTc, resting state toxycist; SmaN, smaller macronuclear nodules. Scale bars: 10 µm.

According to the observation of Feulgen-stained cells (Fig. 8A-C), macronucleus consisting of numerous nodules, quite variable in number, usually between 100 (small-sized individuals) and several hundred: even more than 500 nodules were counted in specimens proximate to cell division on a total of 10 studied cells. Nodules scattered throughout the cell, but less present in anterior and posterior ends (Fig. 8A). In different phases of the ciliate’s life cycle, different types of Ma nodules were observed with different reactivity to Feulgen staining treatment depending on their morphology, sizes, and abundance. Just after cell division, most Ma nodules were densely stained and homogeneously spherical to ovoidal in shape, ∼ 3–4 µm in diameter (Fig. 8B). Before the division process, in most cells two morphologically distinguished subtypes of macronuclear nodules coexisted (Fig. 8C): 1. very abundant, more densely stained ones, showing different shapes, from roundish to irregularly elongated, ∼ 1–2 µm in length (see below for ultrastructural details), which we refer to as “smaller” nodules (Figs. 8C, 9A-C); 2. less brightly stained ones, much less frequently observed (only some tens per cell), almost spherical, ∼ 4–5 µm in diameter (see below for TEM details), we refer to as “larger” nodules (Figs. 8C, 9C). Micronuclei not detectable under the light microscope (neither *in vivo* nor after Feulgen staining) due to their overlapping with the abundant mass of macronuclear nodules (Fig. 8B), but possibly numerous as well; they were observed in TEM treated cells (see below for details; Fig. 9B). One contractile vacuole in posterior cell end (Fig. 7A, D).

**Fig. 9.**
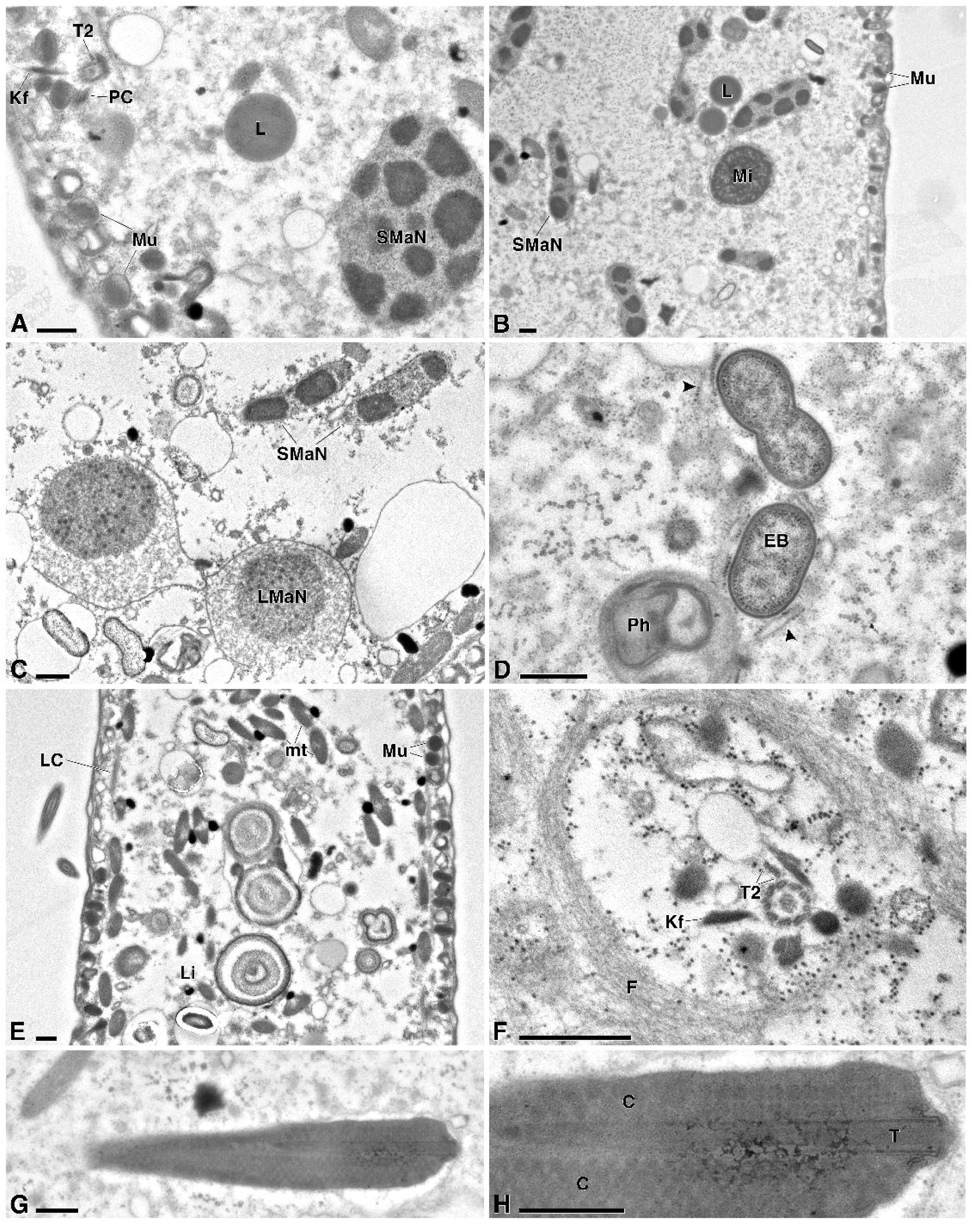
TEM investigation on *Chaenea vermicularis* sp. nov. before cell division. A. Mucocysts inserted in the cortex near two monokinetids of a somatic ciliary row showing part of their typical roots (kinetodesmal fiber and postciliary microtubules) plus the peculiar accessory transverse microtubules T2, and a roundish smaller macronuclear nodule. B. Several elongated smaller macronuclear nodules, including a dividing one, and a micronucleus. C. Two larger macronuclear nodules (characterized by a more electrondense centrally located chromatin core, encircled by a finer homogeneous peripheral chromatin) besides two elongated smaller macronuclear ones. D. Couple of endobacteria (one of which is dividing) close to some rough endoplasmic reticulum cisternae. E. Mucocysts under the cortex and classical mitochondria distributed deeper in the cytoplasm are separated by the *lamina corticalis*; several lithosomes are visible in the cytoplasm. F. A transversely-sectioned somatic monokinetid with part of its roots (kinetodesmal fiber and accessory transverse microtubules T2), encircled by filaments. G, H. Oblique section of a toxicyst at lower (G) and higher (H) magnification to show its layered structure, with an inner tube surrounded by a capsule with a honeycomb-like appearance. Arrowhead indicates rough endoplasmic reticulum cisternae; C, capsule; EB, endobacteria; F, filaments; Kf, kinetodesmal fiber; L, lipid reserve drop; LC, *lamina corticalis*; Li, lithosome; LMaN, larger macronuclear nodules; Mi, micronucleus; mt, mitochondria; Mu, mucocysts; PC, postciliary microtubules; Ph, phagosome; SmaN, smaller macronuclear nodules; T, inner tube; T2, accessory transverse microtubules. Scale bars: 0.5 µm.

In the cortex, numerous mucocysts distributed in regular longitudinal stripes between the ciliary rows (Figs. 7B; 8D, E) (Figs. 7C, 9B, E, see below for ultrastructural details); each strip consists of 5–7 rows of organelles in the equatorial region of the ciliate, while only 4–5 rows at both the anterior and the posterior ends (not shown). Each mucocyst *in vivo* appears as a roundish to oval body, ∼ 0.3 µm in diameter (Fig. 8E). A single type of toxicysts, mace-like in shape, with a tapering posterior end, attached to oral bulge (not shown) and sparse in the cell (Figs. 7A, B). They are ∼ 7 µm long and ∼ 1µm wide in their anterior portion *in vivo* when measured in the resting state (Figs. 7A, B). After the squashing of the cell, among the toxicysts observed *in vivo* there are several extruded ones, recognizable by their longer tip if roughly compared with the resting ones (Figs. 8F, G). Resting state toxicysts were also observed in the cytoplasm in TEM preparations (Fig. 9G, H, see below).

According to *in vivo* and SEM observation, several rows of somatic cilia reach the anterior end of the trunk ending with the circumoral cilia and take an oralized character anteriorly, pointing forward, and including the dorsal brush (Fig. 7A, B E-H). Dorsal brush consisting of four rows, two lateral ones with ∼ 7–12 closely located cilia (SEM length, ∼ 1.8 µm), plus two central rows with 15–20 cilia. Cilia of dorsal brush rows distributed following a zigzag pattern (Fig. 7B, F-H); they are in continuity with somatic cilia (Fig. 7A, B, F-H), which are slightly helically arranged, and roughly 2.0–2.5 times shorter than the latter *in vivo*. Somatic kineties longitudinal, consisting of 8–12 rows of monokinetids (Figs. 7A, B, F, G, I, 8D). Somatic cilia length *in vivo* ∼ 4.4 µm (mean) (SEM length of cilia, ∼ 4.2 µm).

### Notes on the subcellular structure

In the cortical layer, under the plasma membrane, there are flat alveoli and very numerous perpendicularly inserted mucocysts (∼ 0.3 × 0.4 μm) (Figs. 7C, 9A, B, E). In TEM preparations, they show a typical morphology, i.e., a more electron dense inner core with a paracrystalline material distributed along the extrusome length, and a lighter periphery (Fig. 9A, B, E). Under the mucocysts, a *lamina corticalis* is visible lined by numerous classical mitochondria, with tubular cristae, ∼ 1 μm × 0.6 μm in size (Fig. 9E); mitochondria are distributed deeper in the cytoplasm as well.

Monokinetids of the somatic ciliary rows spaced ∼ 3.3 μm apart, each encircled by a dense filamentous layer; monokinetids’ fibrillar associates typical of the litostomateans, i.e., a postciliary ribbon at triplet 9, a laterally directed kinetodesmal fibril at triplets 6 and 7, and the tangential transverse ribbon T1 associated with triplets 3 and 4, plus the so-called “T2”, namely the additional radial transverse microtubules associated with triplet 5 (Fig. 9A, F) (for a reference see Lynn et al. 2008).

In the cytoplasm: 1. Several quite randomly distributed, almost coccoid bacteria (size: ∼ 0.7–0.8 × 0.4–0.5 µm) (Fig. 9D). They are encircled by a clear halo and apparently there is not any symbiosome to separate them from the host cytoplasm; indeed, they are often in direct contact with host’s rough endoplasmic reticulum cisternae. Additionally, they are covered by a double membrane and possess a clearer inner cytoplasm and a ribosomes-rich periphery (Fig. 9D). 2. Medium-sized lipid droplets (∼ 0.75 µm in diameter) (Figs. 7A, B, 9A, B) and lithosomes, i.e., membrane-bound organelles (∼ 4.8 µm in diameter) formed by material laid down in concentric layers (Fig. 9E). 3. The nuclear apparatus: in the cells before the division process, the more abundant smaller macronuclear nodules, appear roundish (∼ 1–2 µm in diameter) to slender and elongated (∼ 1.5 × 0.5 µm in size) in shape (Fig. 9A-C), and the rarer larger nodules are almost spherical (∼ 2.5 µm in diameter) and layered, i.e., with a more electrondense centrally located core of chromatin, encircled by a finer homogeneous peripheral chromatin (Fig. 9C). The smaller macronuclear nodules sometimes appear as two pieces linked by a thin isthmus in between, likely indicating an ongoing division process (Fig. 9B). 4. Rather large, round shaped to ovoid micronuclei (∼ 1.8–2.0 µm in diameter), containing a dense chromatin (Fig. 9B). 5. Small phagosomes (∼ 0.8 µm in diameter) with a digested content (Fig. 9D). 6. Toxicysts in the resting state, at least∼ 6.5 μm long, but often their whole length is not appreciable in TEM sectioned cells due to oblique sectioning; they are ∼1µm wide as measured in their anterior portion (Fig. 9G, H). Toxicysts structure typical, i.e., including an inner tube (∼ 0.2 µm in diameter) surrounded by an outer tube (the capsule), but the latter manifests an uncommon honeycomb-like appearance due to the presence of alternating zones of different electron density; they are generally encircled by a membrane, with a reduced space between the latter and the capsule, sometimes showing a finely filamentous content filling the space around (Fig. 9G, H). Sometimes, developing toxicysts were detected in the cytoplasm as well; their “juvenile” state was clear given the presence of a larger space between their membrane and the vacuole membrane (not shown).

### Occurrence and ecology

During its permanence in the laboratory, the *C. vermicularis* population grew by predating on unidentified smaller protists of the original sample, which are compatible with the observed small phagosomes (see above and Fig. 9D) likely indicating a very fast digestion process In the laboratory, the ciliate mainly crawled over the substratum. Cyst formation was not observed.

### Gene sequence

The 18S rDNA sequence of *C. vermicularis* obtained from PCR resulted 1,598 bp long, and it has been deposited in NCBI GenBank database with the accession number PQ628062. The 18S rDNA sequence of *C. vermicularis* showed the highest identity with the sequence of *Trachelophyllum* sp. (JF263452): 90.37% (20 gaps, 135 mismatches). The identity value with *Chaenea* sp. (MF474337) resulted 89.34% (25 gaps, 144 mismatches). Identity values with other *Chaenea* sequences ranged from 86.43% to 87.45%.

### Mitochondrial genome

The mitochondrial genome of *C. vermicularis,* resulted in a single linear molecule of 40,677 bp (for further details see the “Mitochondrial genome” section, below).

### Microbial consortium

The screening of the preliminary assembly for bacterial 16S rDNA sequences revealed a single bacterial species (OTU #12) putatively associated with *C. vermicularis.* The retrieved 16S rDNA sequence belongs to an uncultured *Cytophagia* (*Bacteroidota*) closely related to “*Candidatus* Amoebophilus”, OTU #12. Details about sequence’s length, coverage, accession number, and best BLAST hits are given in Table 2. As for the ultrastructural details, see above.

### Phylogenetic analyses (FIG. 10)

The 18S rDNA-based phylogenetic analysis placed the sequences of *L. venatrix* and *P. famelica* inside a group formed by sequences of other members of *Lacrymaria* and *Phialina* genera. The sequence of *L. venatrix* branched basally from a group formed by the cluster of *Phialina* sequences (containing also the sequence of *P. famelica*), and a cluster of sequences comprising other *Lacrymaria* sequences (e.g., *L. olor* MN030556, *L. maurea* MF474344 and *L. marina* MF474343), while the sequence of *P. famelica* is sister of two sequences of *P. salinarum* (EU242508 and FJ876975) with high statistical support (1.00, 93%). Our analysis failed to recover neither *Lacrymaria* nor *Phialina* as monophyletic group, coherently with previous phylogenetic analyses (Vďačný et al. 2014; Huang et al. 2018; Tang et al. 2023). The sequence of *C. vermicularis*, instead, clustered in a well-supported (1.00, 96%) monophyletic group formed by all the other sequences of representatives of the genus. The sequence of *C. vermicularis* branches early compared to all the other sequences of *Chaenea* species; it is connected to them by a rather long branch mirroring a higher variability in its sequence compared to the other members of the genus (18S rDNA identity values ranging from 85.59% to 89.34%).

### Mitochondrial genome (FIG. 11)

The three reconstructed mitochondrial genomes resulted in each case in a single linear molecule 62,315 bp, 52,439 bp, and 40,677 bp long for *Lacrymaria venatrix*, *Phialina famelica,* and *Chaenea vermicularis*, respectively. The gene content of the three mitochondrial genomes is overall fairly similar, composed of ribosomal proteins, proteins for cytochrome c oxidases, cytochrome c reductase or NADH dehydrogenase subunits, and transfer and ribosomal RNAs (Supplementary Table 1). Among these proteins of known functions, no new proteins (i.e., not already identified in other ciliates’ mitochondrial genomes) were found. However, among the several predicted proteins with unknown function identified in the three genomes, six showed a significant similarity in the three litostomatean mitochondrial genomes, while not having recognizable homologous in other available ciliate mitochondrial genomes, suggesting a set of homologous proteins unique of this lineage of ciliates.

As expected, *L. venatrix* and *P. famelica* showed a higher degree of synteny compared to *C. vermicularis,* even if a chromosome rearrangement seems to have occurred in *Lacrymaria* mitochondrial genome’s terminal part (Fig. 11).

## DISCUSSION

### Issues in the systematics of the *Lacrymaria* and *Phialina* genera

The taxonomic history of the genera *Lacrymaria* and *Phialina* is quite complex. Originally, they were both included (*Lacrymaria* under the name “*Lacrimatoria*”) by Jean Baptiste Bory de Saint-Vincent in his *Histoire naturelle des zoophytes* (1824, also see Rajter et al., 2019) in the class “Microscopica”, although in two different orders (respectively “Gymnodes” and “Trichodes”). At that time, “*Lacrimatoria*” and *Phialina* included species described earlier by Otto Müller (1773, 1786) as *Vibrio olor* and *Trichoda pupula* (now referred to as *Lacrymaria olor* and *Phialina pupula*, respectively). It is worth noting that contemporary taxonomic categories often do not correspond to modern ones: for example, in the *Histoire naturelle* the bacterial genus *Vibrio*, from which Bory de Saint-Vincent had removed *L. olor*, also includes species of nematodes.

Both genera were described as having a contractile neck, with a main discriminant of the position of the cytostome: “*Lacrimatoria*”, later on changed in *Lacrymaria* by Ehrenberg (1831), having a terminal one, *Phialina* a lateral one. Under these criteria new species were described, though the so called “holotrichous” genera remained rather labile, as authors such as Ehrenberg (1833) and Kent (1882) frequently moved species into or out genera such as in the case of *Amphileptus* and *Trachelocerca*.

However, later progress in microscopy demonstrated that both *Lacrymaria* and *Phialina* have in fact terminal cytostomes. Alfred Kahl (1930) formally abolished the distinction by collapsing *Phialina* within the genus *Lacrymaria*. With re-descriptions and synonymizations, he also brought down to 13 the number of *Lacrymaria* species, most of which are still valid ones. Consequently, the species expanded again, now totalling approximately 68 presumably valid species. The largest pulse of descriptions occurred between 1955 and 1965, owed to Jean Dragesco and A. L. Vuxanovici; many of these species, however, are infrequently cited in literature.

When Foissner (1983) studied *Lacrymaria* species with new staining techniques, he found it useful to introduce a new discriminant: he redescribed *Lacrymaria* including a long contractile neck as a defining feature and re-established the genus *Phialina* for the related species lacking it. That is the currently accepted standard, which later descriptions have been based on.

But this might be changing once again in the future. Indeed, previous 18S rDNA-based molecular phylogenies (e.g.,Vďačný et al., 2014; Huang et al., 2018; Wang et al., 2019; Tang et al., 2023) and the present work show that the genera *Lacrymaria* and *Phialina* are nested within each other, that neither, on its own, is monophyletic, and that the set of both, were they to be combined into a single genus, would indeed be monophyletic (Fig. 10). Considering that the name *Lacrymaria* has been used by far the most, and was indeed the chosen by Kahl (1930) when he combined the two genera, the retaining of that name in the event of any further genus fusion would presumably serve better the purpose of the Principle of Priority which, according to the ICZN (23.2), “is to be used to promote stability and it is not intended to be used to upset a long-accepted name in its accustomed meaning” (ICZN, 1999).

**Fig. 10.**
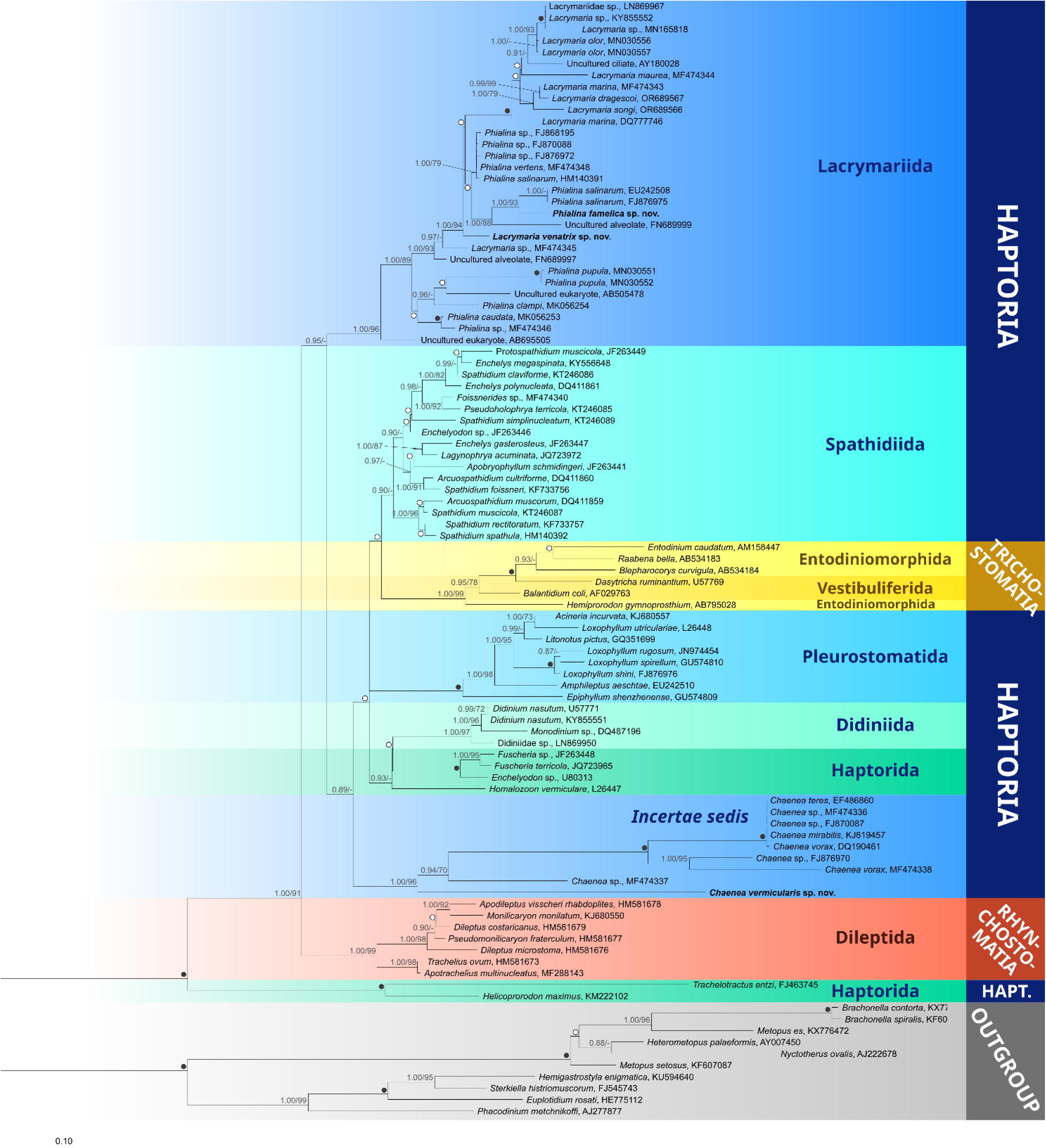
The 18S rDNA-based phylogenetic tree of Class Litostomatea showing the position of the three studied species, i.e., *Lacrymaria venatrix* sp. nov. (accession number: PQ628060), *Phialina famelica* sp. nov. (accession number: PQ628061), and *Chaenea vermicularis* sp. nov. (accession number: PQ628062). Numbers associated to nodes represent posterior probability from Bayesian inference (BI) and bootstrap value from maximum likelyhood (ML) analyses, respectively (only values of BI > 0.85 and ML > 70% are shown). *Black dots* represent the highest statistical support (BI = 1.00 and ML = 100); *white dots* indicate non-significant statistical support (BI < 0.85 and ML < 70%). Sequences obtained in the present work are in bold.

**Fig. 11.**
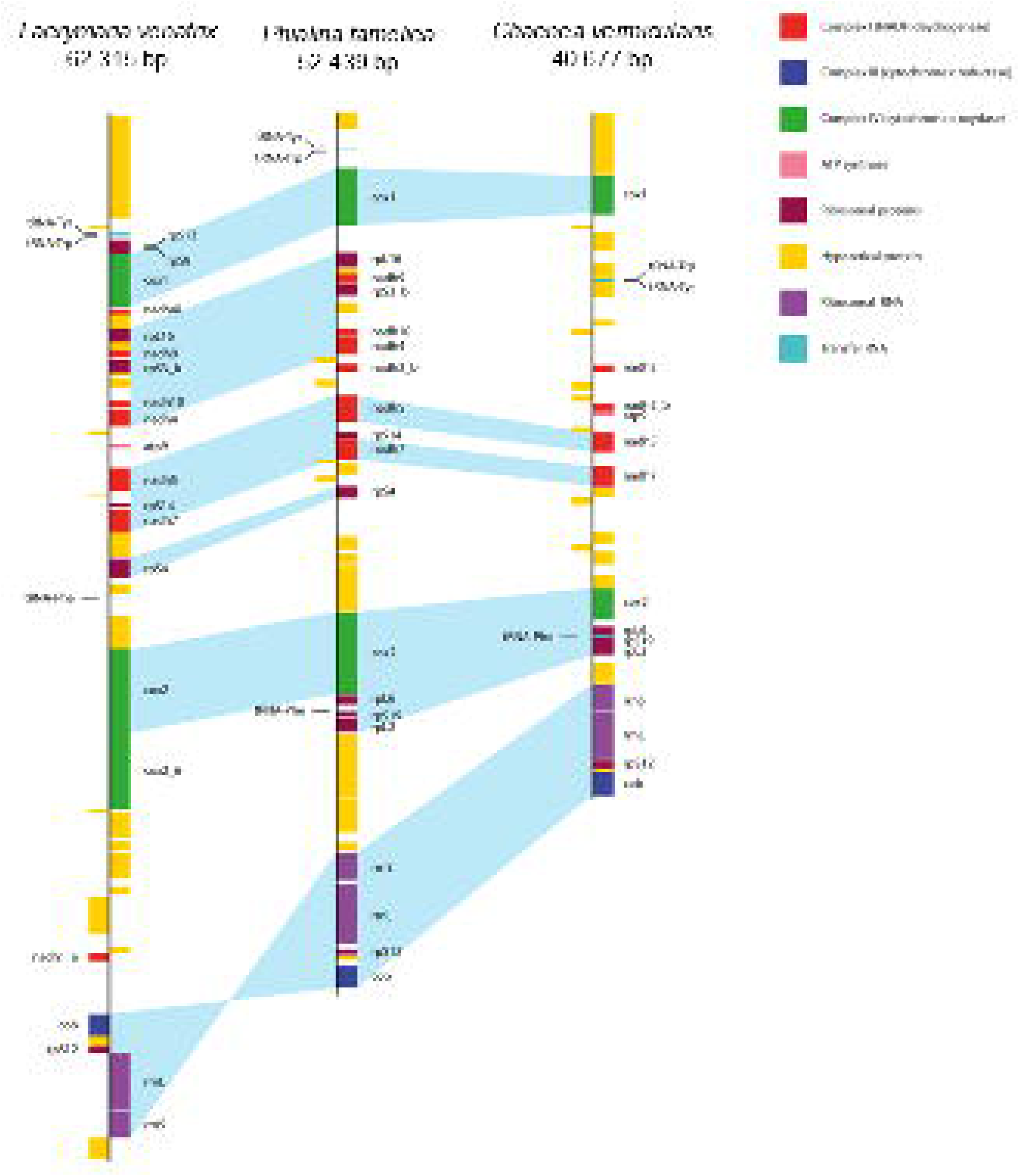
Gene map of the recovered mitochondrial genomes of *Lacrymaria venatrix* sp. nov., *Phialina famelica* sp. nov., and *Chaenea vermicularis* sp. nov., and their comparison in terms of gene order. Syntenic regions are highlighted by light blue connectors across the three maps, including those with conserved gene order but inverted orientation.

Although we are aware that probably, in the future, the current systematics of *Lacrymaria* and *Phialina* will be critically reviewed and hopefully fixed, since a systematic review is not the goal of the present paper, we herein describe our novel species according to the currently accepted standard which refers to Foissner (1983).

### *Lacrymaria venatrix* and *Phialina famelica* are novel species

From the standpoint of morphology, *L. venatrix* sp. nov. does not exhibit individually unique features, but rather a unique, novel combination of features typical of *Lacrymaria* species. Therefore, based on morphological relevant traits, the ciliate should be compared to *L. olor* (Foissner, 1997), *L. marina* (Song and Packroff, 1997; Dong et al., 2010), and *L. songi* (Tang et al., 2023) (Table 5).

**Table 5.** Comparison of *Lacrymaria venatrix* sp. nov. with selected similar congenerics.

| Character | *L. olor* | *L. marina* | *L. songi* | *L. venatrix* |
| --- | --- | --- | --- | --- |
| Habitat | Freshwater | Marine | Marine | Freshwater |
| Body length (Min-Max (Mean)) | 118–220 (152) | 133–171 (149) | 88–187 (130) | 73–121 (92) |
| Body width (Min-Max (Mean)) | 15–24 (17) | 4–39(34) | 15–39 (26) | 14–23 (19) |
| Neck: total length ratio | Up to 4:5 | Up to 2:3 | ~1:2 | Up to 1:2 |
| Somatic kineties, number | 13–16 (14) | 16–23 | 12–16 | 17–18 |
| Monokinetids per kinety, number | ~90–200 | -- | 80–180 | -- |
| Mucocysts, size (IV) | 0.4 × 0.6 | 0.2–0.5 (diam) | 0.4 (diam) | ~0.6 (diam) |
| Rows of Mu per strip, number | ~8 | 4–5 | 5–6 | 5–6 |
| Extrusomes (number types, type, length) | 2*, Tc, ~11/ ~2.5 | 1, Tc<sup>□</sup>, ~10 | 2, Tc<sup>□</sup>, ~10/ ~4 | 1, Tc, ~12 |
| Macronuclear nodules (number, shape) | 2, ellipsoidal | 1, ellipsoidal | 2, irregular | 2, ellipsoidal |
| Macronucleus length | 9–14 (11) | 29–46 | 10–32 (18) | 18–36 (28) |
| Macronucleus width | 8–11 (9) | 11–19 | 6–15 (10) | 8–15 (12) |
| Micronuclei, number | 1 | 1 | 2 | 1 |
| Contractile vacuole (number, place) | 2, lateral | 1, terminal | 1, terminal | 1, terminal |
| Reference | Foissner, 1997 | Song and Packroff, 1997;<br>Dong et al., 2010 | Tang et al., 2023 | Present study |

Measurements in $\mu\text{m}$ . All the measurements are from Protargol stained specimens with retracted neck. diam, diameter; IV, *in vivo*; Max, maximum value; Mean, mean value; Min, minimum value; Mu, mucocysts (including when referred to as “cortical granules” in the paper; see text for details); Tc, toxicysts; --, data not available; \* observed/measured/calculated on the drawing from protargol stained cells on the original paper; <sup>□</sup> the authors did not refer to the observed extrusome (or, in case of more than one type of extrusome, to the longer extrusome type) as “toxicysts”, but based on their appearance, they are called toxicysts here. Marine environment indicates salinity $\geq 30\text{‰}$ .

The size of the trunk of *L. venatrix* (92 × 19 μm) is considerably smaller than in *L. olor* (152 × 17 μm) (Foissner, 1997), *L. marina* (139 × 34 μm) (Song and Packroff, 1997; Dong et al., 2010) or *L. songi* (130 × 26 μm) (Tang et al. 2023). The extended neck-to-trunk ratio in *L. venatrix* (roughly 1:1) is instead closest to *L. marina* (2:1) (Song and Packroff, 1997; Dong et al., 2010).The macronucleus formed by two ellipsoidal nodules is a feature shared by *L. venatrix* and the long-necked *L. olor* (Foissner, 1997), whereas *L. marina* has only a single, ellipsoidal nodule (Song and Packroff, 1997; Dong et al., 2010). A single terminal contractile vacuole, on the other hand, is a common trait of *Lacrymaria* species, but *L. olor* has two (Foissner, 1997). The number of somatic kineties of *L. venatrix* (17-18) is higher than in *L. olor* (13-16) (Foissner, 1997) though still lower than in larger *Lacrymaria* species, such as *L. marina* (16-23). Mucocysts of *L. venatrix* also appear in line with those of the similar *Lacrymaria* species (Table 5), somewhat more densely packed than those of *L. marina* (Song and Packroff, 1997; Dong et al., 2010) (5–6 vs. 4–5 rows) but less packed than those of *L. olor* (Foissner, 1997) (5-6 vs. 8 rows).

However, in terms of overall morphology, *L. venatrix* is rather similar to *L. songi* recently described from brackish water (Tang et al., 2023) (Table 5), though with the following divergent traits. It is significantly smaller in size (73–121 vs. 150–300 µm), with a proportionately bigger macronucleus (28 × 12 vs. 18 × 10 μm) whose nodules are not clearly separated (vs. with an evident gap in between in *L. songi*), and with only one micronucleus (vs. two micronuclei in *L. songi*). The kineties of *L. venatrix* are more numerous (17-18 vs. 12-16). As for the extrusomes: 1. the trunk of *L. songi* contains elongated organelles (toxicysts) of two kinds, the longer often arranged in bundles, diffused in the cytoplasm, and inserted into the oral bulge, and the shorter, only scattered in the main body vs. in *L. venatrix* we observed only one type localized in both cell districts; and 2. In *L. songi* two types of “cortical granules” are reported, one of the two arranged in rows between kineties vs. in our species there is only one type, i.e., mucocysts according to our TEM observation and literature definition (Hausmann, 1978; Rosati and Modeo, 2003), with the same arrangement. In this regard, considering that mucocysts referred to as “cortical granules” are reported in about all the literature concerning the genera object of the present study (i.e., *Lacrymaria, Phialina,* and *Chaenea*), we take this opportunity to stress that species description relying only on the light microscope observation can often provide solely a general categorization (if not a miscategorization) of certain structures, which sometimes can largely affect species characterization and identification. A reliable setting is only possible conducting a subcellular investigation (for example by means of TEM) in parallel to the light microscope one. The predatory behavior of *L. venatrix* does not appear substantially different from that of *L. olor*, which is extensively described in literature (e.g., Fenchel, 1968, Coyle et al., 2019): although the neck of *L. venatrix* is not as lengthy as that of *L. olor*, it is still sufficiently flexible to cover a similar range around the cell’s trunk and reducing the need to move it during predation. Moreover, the mechanism of extension and retraction is likely the same, consisting in a reallocation of cytoskeleton to/from the neck and a remodelling within the neck of elements of the cortical cytoskeleton (Coyle et al., 2019, Banavar et al., 2022; Qin et al., 2024). According to our phylogenetic analysis, *L. venatrix* was embedded within the *Lacrymaria/Phialina* clade, although it did not cluster closely with any other sequences currently available in online databases (Fig. 10), supporting its designation as a novel species.

In the case of *P. famelica*, the peculiar presence of three different kinds of toxicysts inserted in the oral bulge, disclosed by our TEM investigation, is *per se* a distinctive feature of a novel species. Nevertheless, based on morphological relevant traits, the ciliate should be compared to *P. salinarum* (Song et al., 2003), *P. serranoi* (Foissner, 2016), *P. vertens* (Foissner, 1983; Song and Wilbert, 1989), *P. pupula* (Rajter et al., 2019), and *P. vermicularis* (Foissner, 1983; Alekperov, 2005) (Table 6). Foissner et al. (2002) mentioned that *P. minima* is not contractile while *P. famelica* is moderately contractile; thus, the former species is not elaborated upon in this paper. *Phialina serranoi* (Foissner, 2016) and *P. salinarum* (Song et al., 2003) are the species closest to *P. famelica* according to size and general morphology. The latter two cluster together in the phylogeny as well (Fig. 10), even though *P. salinarum* is a marine species; however, this is not surprising as *P. famelica* also originates from an environment with moderate to high salinity, although anthropogenic. However, the number of somatic kineties in *P. famelica* is lower than in *P. salinarum* (mean: 14 vs. 18) and is closer to that of *P. pupula* (Rajter et al., 2019) and *P. serranoi* (Foissner, 2016), though falling short of other species of similar size such as *P. vertens* (Foissner, 1983; Song and Wilbert, 1989), and *P. vermicularis* (Foissner, 1983; Alekperov, 2005). On the other hand, the number of monokinetids per kinety (mean: 55) in *P. famelica* is much higher than in all other considered species: indeed, no other species seem to reach past 40.

**Table 6.** Comparison of *Phialina famelica* sp. nov. with selected similar congenerics.

| Character | *P. salinarum* | *P. serranoi* | *P. vertens* | *P. pupula* | *P. vermicularis* | *P. famelica* |
| --- | --- | --- | --- | --- | --- | --- |
| Habitat | Marine or brackish | Freshwater (sludge) | Freshwater | Freshwater | Freshwater | Brackish (sludge) |
| Body length (Min-Max (Mean)) | 60–100 IV | 58–87 (72) P | 50–85 | 53–115 (75) | 80–100 µm (German pop.); 40–60 (Austrian pop.); 110–130 (Azerbaijani pop.) IV | 35–66(48) P |
| Body width (Min-Max (Mean)) | 15–30 IV | 18–27 (23) P | -- | 17–42 (26) | -- | 15–34 (23) P |
| Somatic kineties (n) | 17–21 | 14–16 | 15–20 | 12–16 (15) | 18–20 | 12–16 (14) |
| Monokinetids per kinety | ~33 | 17–35 (25) | ~30–40* | 13–33 (22) | 35* | 33–68 (55) |
| Mucocysts, size (IV) | 0.5–1.0 (diam) | 0.4 × 0.8 | -- | 0.4 × 0.8 | -- | 0.2–0.4 (diam) |
| Rows of Mu per strip, number | 4–5 | -- | -- | 7–8 | -- | ~5 |
| Extrusomes (number types, type, length) | 1, Tc<sup>□</sup>, ~10 | 2, Tc<sup>□</sup>, ~12/~3 IV | 1, Tc<sup>#</sup>, ~12* | 1, Tc<sup>□</sup>, 10 IV | 1, Tc<sup>#</sup>, ~15** | 1, Tc, ~8.5 P |
| Special cytoplasmic inclusions | absent | Absent | absent | highly refractive dumbbell-shaped inclusions | absent | refractive inclusions |
| Macronuclear nodules (number, shape) | 1, ellipsoidal | 1, ellipsoidal or reniform | 1, ellipsoidal or dumbbell | 1, ellipsoidal | 1, ellipsoidal to reniform | 1, ellipsoidal or, more frequently, reniform |
| Macronucleus length | 12–21 | 9–17 (14) | 15–20 | 10–20 (15) | 15 (Austrian pop.) P | 17–38 (27) |
| Macronucleus width | 5–9 | 5–12 (9) | 15 | 5–16 (10) | 10 (Austrian pop.) P | 7–15 (10) |
| Micronuclei, number | 1–2 | 1 | 1 | 1 | -- | 1 |
| Contractile vacuole (number, place) | 1, terminal | 1, lateral | 1, terminal | 1, subterminal | 1, subterminal or terminal | 1, terminal |
| Reference | Song et al., 2003, Dong et al., 2010 | Foissner, 2016 | Foissner, 1983; Song and Wilbert, 1989 | Rajter et al., 2019 | Foissner 1983; Alekperov 2005 | Present study |

Measurements in µm. diam, diameter; IV, *in vivo*; Max, maximum value; Mean, mean value; Min, minimum value; Mu, mucocysts (including when referred to as “cortical granules” in the paper; see text for details); n, number; P, protargol staining; pop., population; --, data not available. <sup>□</sup> the authors did not refer to the observed extrusomes as “toxicysts”, but based on their appearance, they are called toxicysts here. <sup>#</sup>, author indicated as “protrichocysts” the extrusomes, but based on their appearance, they are called toxicysts here; \* observed/measured/calculated on the drawing based on protargol stained cells on the original paper. \*\* observed/measured/calculated on the drawing/micrographs from *in vivo* cells on the original paper. Note that, according to our TEM data, *P. famelica* bears three different types of toxicysts inserted in its oral region (see main text for details). Marine environment indicates salinity ≥ 30‰.

The angle at which the kineties of *P. famelica* rest when the body is contracted, is much higher than in all the abovementioned species being *P. salinarum* (Song et al., 2003) the closest (∼ 45 vs. ∼ 30°); however, this difference might be explained simply by a different (greater) state of contraction. *Phialina serranoi* is another sewage species reported by Foissner (2016) in an Austrian WWTP. It has a contractile vacuole located far subterminal, so very unusual in the *Phialina* species (*P. famelica* included). Moreover, *P. serranoi* can also be distinguished from our species by having larger-sized protargol-stained cells (72 × 23 vs. 48 × 23 μm), a smaller-sized macronucleus (14 × 9 vs. 27 × 10 μm) and a lower number of monokinetids per somatic kinety (mean: 25 vs. 55). Finally, *P. famelica* can be separated from *P. vertens* (Foissner, 1983; Song and Wilbert, 1989) and *P. vermicularis* (Foissner, 1983; Alekperov, 2005) in having less somatic kineties (mean: 14 vs. 15–20 and 18–20, respectively), more monokinetids per each somatic kinety (mean: 55 vs. 30–40 and 35, respectively), and a higher, although still moderate, contractility (30–50% vs. 12–16% and 16–25%, respectively).

To date, molecular data are available for *L. dragescoi*, *L. olor*, *L. marina*, *L. maurea*, *L. songi*, and for *P. caudata*, *P. clampi*, *P. famelica*, *P. pupula*, *P. salinarum*, plus *L. venatrix* and *P. vertens* (Fig. 10); note that after a careful check of the published sequences some doubts should be expressed about the identification of MF474348 (*P. vertens*), mainly because it was collected from Jiaozhou Bay, China, which is a marine habitat (Liao, 2021). From a phylogenetic perspective, *L. venatrix* fell basally to the *Lacrymaria* + *Phialina vertens* + *Phialina salinarum* – clade and *P. famelica* as sister of *P. salinarum* (EU242508, FJ876975) (Fig. 10). These results are in line with those of other studies (e.g., Vďačný et al., 2014; Huang et al., 2018; Wang et al., 2019; Tang et al., 2023), underlining the paraphyly of the two genera. However, it is worth noting that the basal position of *L. venatrix* may suggest that the extensible neck could be a morphological feature acquired (or lost?) independently many times, during the evolution in this clade.

In conclusion, based on the combination of morphological traits, 18S rDNA sequence identity, and the phylogeny of *L. venatrix* and *P. famelica*, we confidently propose these two as novel species. Hower, as mentioned above, we are aware that a systematic review of *Phialina* and *Lacrymaria* genera is needed and that, possibly, in the next future one of the two novel species could be renamed according to a new classification.

### *Chaenea vermicularis* is a novel species

Considering both habitat and morphological features (such as cell contractility, body length, length-to-width ratio, somatic kineties rows number, mucocysts arrangement, extrusomes other than mucocysts, and number of macronuclear nodules), *C. vermicularis* should be compared with *C. teres* (Petz et al., 1995), *C. vorax* (Song and Packroff, 1997), *C. paucistriata* (Fan et al., 2015), and *C. mirabilis* (Kwon et al., 2014), which are marine representatives of the genus (Table 7). Unfortunately, not all the morphological data available in the literature for these *Chaenea* species (Petz et al., 1995; Song and Packroff, 1997; Kwon et al., 2014; Fan et al., 2015) can be considered exhaustive; an emblematic case of this situation is the rather poor morphological description of *C. vorax* (Song and Packroff, 1997). In particular, the morphological descriptions of the four species are far from sufficiently clear regarding fundamental aspects such as the features and the arrangement of toxicysts and mucocysts. Additionally, in *C. teres* and *C. paucistriata* “cortical granules” are reported (Petz et al., 1995; Fan et al., 2015) instead of mucocysts. As anticipated in the previous section, according to our ultrastructural analysis *C. vermicularis* possesses typical mucocysts (Hausmann, 1978; Rosati and Modeo, 2003) like *L. venatrix* and *P. famelica*; this finding supports previous TEM observations by Lipscomb and Riordan (1990) in *C. teres*, and by Fauré-Fremiet and Ganier (1969a) in *C. vorax*, where “corps mucifères” were described. Therefore, we will consider cortical granules of *C. teres* and *C. paucistriata* as mucocysts as well. However, *C. vorax sensu* Song and Packroff (1997) and *C. mirabilis* apparently diverge, given that numerous large granules, approximately 3-8 μm in size, are only described in the endoplasm (Song and Packroff, 1997), which are not to be considered as cortical granules in the former (lipid droplets?), and no cortical granules can be discerned in the cortex of the latter, where only 3–5 special highly refractive subcortical globules are present near the dorsal brush rows (Kwon et al., 2014).

**Table 7.** Comparison of *Chaenea vermicularis* sp. nov. with selected similar congenerics.

| Character | *C. teres* | *C. vorax* | *C. paucistriata* | *C. mirabilis* | *C. vermicularis* |
| --- | --- | --- | --- | --- | --- |
| Habitat | Marine | Marine | Marine | Marine | Marine |
| Contractility | Moderate | -- | High | High | Moderate |
| Body length (Min-Max (Mean)) | 120–170 (195‡) | 100–180 (140‡) | 180–250 (215‡) | 60–100 (75‡) | 130–520 (320) |
| Length/Width ratio (IV) | 8.3:1** | 8.4:1** | 12–14:1 | 7:1 | 23:1 |
| Somatic kineties, number | 12 § | 11–12 | 8 | 12 | 8–12 |
| Mucocysts, size (IV) | -- | absent | < 0.5 (diam) IV | absent | ~ 0.3 (diam) |
| Rows of Mu per strip, number | 3–4** | -- | 3–4** | N/A | 5–7 |
| Extrusomes (number types, type, length) | 1, Tc, 11* | 1, Tc, 6**, 5–6 P | 1, Tc, 8 IV | 2<sup>□</sup>, Tc, 6–8 IV, 3 P | 1, Tc, 6.6–7 IV |
| Macronuclear nodules (n) | many* | 80–110 | 63–94 | 11–21 | hundreds |
| Reference | Petz et al., 1995 | Song and Packroff, 1997 | Fan et al., 2015 | Kwon et al., 2014 | Present study |

Measurements in $\mu\text{m}$ . IV, *in vivo*; Ma, macronucleus; Max, maximum value; Min, minimum value; Mu, mucocysts (including when referred to as “cortical granules” in the paper; see text for details); N/A, character non present; P, protargol staining; SK, somatic kineties; Tc, toxicysts; --, data not available; ‡ mean calculated on Max and Min values of the original paper; \* observed/measured/calculated on the drawing from protargol stained cells on the original paper; \*\* observed/measured/calculated on the drawing/micrographs from *in vivo* cells on the original paper; §, mean value; □, the authors describe two types of extrusomes: the longer type is likely a form of toxicyst, so it is referred to as such here, even though the authors did not categorize it this way (the second type has a completely different morphology, thus it has not been considered here). Marine environment indicates salinity $\geq 30\%$ . Note that according to Song and Packroff (1997) numerous granules, approximately 3–8 $\mu\text{m}$ in size are present in the endoplasm, thus cortical granules are considered absent.

*Chaenea vermicularis* can be distinguished from all four mentioned congenerics based on the different traits listed below (Table 7). 1. Cell contractility: slightly contractile vs. *C. teres* (Petz et al., 1995), only slightly contractile cell; *C. vorax*, no data (Song and Packroff, 1997); *C. paucistriata* (Fan et al., 2015) and *C. mirabilis* (Kwon et al., 2014), very flexible and contractile cell. 2. Body length *in vivo* (mean): 320 µm vs. *C. teres*, 195 µm; *C. vorax*, 140 µm; *C. paucistriata*, 215 µm; *C. mirabilis,* 75 µm. 3. Cell length-to-width ratio *in vivo*: 23:1 vs. *C. teres*, 8.3:1; *C. vorax*, 8.4:1; *C. paucistriata*, 12-14:1; *C. mirabilis*, 7:1. 4. Number of somatic kineties: 8–12 vs. *C. teres*, 12; *C. vorax*, 11–12; *C. paucistriata*, 8; *C. mirabilis*, 12. 5. Number of mucocysts rows per strip: 5–7 vs. *C. teres*, 3–4; *C. vorax*, no available data; *C. paucistriata*, 3–4; *C. mirabilis*, no available data. 6. Number of macronuclear nodules: several hundred vs. *C. teres*, “many”; *C. vorax*, 80–110; *C. paucistriata*, 63–94; *C. mirabilis*, 11–21.

Concerning extrusomes other than mucocysts, notably, all of these *Chaenea* species possess only one type, namely toxicysts, except *C. mirabilis* which shows two types, a longer one, rod-shaped, which is not explicitly referred to as toxicysts by the Authors but can be very likely identified as such based on its morphological features and localization in the cell, plus a second type, referred to as “narrowly to broadly teardrop-shaped” (Kwon et al., 2014), which has not been considered for the purposes of morphological comparison (Table 7). Resting state toxicysts length *in vivo* for all the other species of Table 7 is similar to that measured in *C. vermicularis*: 6.6–7 µm vs. *C. vorax*, 6 µm; *C. paucistriata*, 8 µm; *C. mirabilis*, 6–8 µm.

Additionally, *C. teres*, *C. vorax*, and *C. paucistriata* did not manifest the presence of two subtypes of macronuclear nodules such as those distinctively present in our species during part of the ciliate life cycle (Petz et al., 1995; Song and Packroff, 1997; Fan et al., 2015). On the contrary, in the case of *C. mirabilis* such a remark is found concerning the macronucleus: “It consists of 11– 21 doughnut-shaped or sometimes horseshoe shaped macronuclear nodules” (Kwon et al., 2014). As for *C. vermicularis*, the Feulgen-staining procedure supported by TEM observation clearly disclosed the concurrent presence of smaller irregular macronuclear nodules and larger spherical ones with different abundance (Fig. 8A-C, 9A-C) exclusively in ciliates proximate to cell division. Although we did not specifically investigate the dynamics of the macronuclear variety in *C*. *vermicularis*, to explain the nodules heterogeneity we hypothesize that, possibly, some peculiar process takes place in the macronuclear nodules before cell division, namely some of them fuse (or develop) in a larger form that is able to divide, while, at the same time, the most numerous, smaller and irregular ones elongate and split into two or disappear to develop again later, at some stage after the cell division. Indeed, the presence of macronuclear nodules that elongate and divide before cell division in *C. vermicularis* supports the observation on *C. vorax* by Fauré-Fremiet and Ganier (1969a), while these Authors excluded any fusion of nodules in their species.

Only three ultrastructural studies have been previously performed on *Chaenea* spp.: two of them deal with *C. vorax* (Fauré-Fremiet and Ganier, 1969a, b), while the third deals with *C. teres* (Lipscomb and Riordan, 1990). However, the species described by Lipscomb and Riordan (1990) shows only three rows of clavate cilia in the dorsal brush. The somatic monokinetids’ roots of *C. vermicularis* appear typical of litostomatean predators (Lynn, 2008), and so are the layered toxicysts and the mucocysts according to their general ultrastructure (Hausmann, 1978; Rosati and Modeo, 2003) confirming previous observations on *Chaenea* species (Fauré-Fremiet and Ganier, 1969a; Lipscomb and Riordan, 1990). As a peculiarity, we could observe in *C. vermicularis* the presence of a *lamina corticalis*, the ecto-endoplasmic fibrillar layer typical of the litostomes (Lynn, 2008), which has not been observed neither in *C. vorax* (Fauré-Fremiet and Ganier, 1969a) nor in *C. teres* (Lipscomb and Riordan, 1990). Finally, in the latter two species, lithosomes, frequently found in TEM sections of spirotrichean ciliates such as hypotrichs and stichotrichs (Lynn, 2008), were not observed, while they were obvious in the cytoplasm of *C. vermicularis*, reminding of those recorded in the predator *Helicoprorodon multinucleatum* (Lipscomb and Riordan, 1991).

Molecular data are available only for three out of the four abovementioned species of *Chaenea*, i.e., *C. vorax*, *C. teres*, and *C. mirabilis*, plus some *Chaenea* spp. In general, the *Chaenea* clade is presently under debate, since Zhang and colleagues (2024) hypothesized it might represent a higher-level taxon within the Subclass Haptoria, leaving the genus as *incertae sedis*. In the present study we followed their interpretation.

Moreover, phylogenetic reconstruction based on 18S rDNA gene sequence indicates that *C. vermicularis* split off basally respect to all the other genus members, with which it showed relatively low identity values (∼ 85 to 89%). The significant genetic differences observed between *C. vermicularis* and other related species might even indicate that it could represent a new genus. However, we chose to take a cautious approach, as this topic warrants further investigation. In this context, we believe that our multidisciplinary characterization of *C. vermicularis*, which includes subcellular analysis using transmission electron microscopy, may help a possible reevaluation of the genus’ status in the future.

In conclusion, what can be at present surely inferred based on the whole body of data (morphological plus molecular ones), is that our *Chaenea* species has to be considered new to science.

### Mitochondrial genomes

In the present paper, we report the first annotation of the mitogenome of representatives of the Class Litostomatea. The overall length and content of mitogenomes of *L. venatrix, P. famelica* and *C. vermicularis* resulted in line with other ciliates’ mitochondrial genomes (Swart et al., 2012; Serra et al., 2020, 2021). It is also worth to notice the presence of several proteins in the mitochondrial genome which seems to be unique of this lineage of ciliates, and therefore potentially related to their predatory lifestyle. It is also curious to notice that their mitogenomes showed a lower level of synteny in respect to other representatives of other classes, e.g. Spirotrichaea or Oligohymenophorea (Liao et al., 2021; Serra et al., 2021; Xu et al., 2025), with fewer blocks of genes whose order seems conserved among the three genomes at class level.

### Microbial consortium associated with the three predators

In the cytoplasm of all the three studied litostomateans the presence of bacteria was observed by TEM analysis, and also from molecular screening several 16S rDNA sequences were retrieved from the ciliates’ assemblies (Table 2). However, unfortunately, due to the disappearance of *P. famelica* and *C. vermicularis* populations, FISH analyses could only be performed on *L. venatrix*, although results did not show any clear signal of the retrieved OTUs (#1–4) in the cytoplasm of *Lacrymaria* specimens.

The OTU #1 (Table 2) retrieved in *Lacrymaria,* initially, seemed the most probable candidate to match the bacteria observed in its cytoplasm by means of TEM, being *Bdellovibrio* well known as an obligate intracellular organism (Cavallo et al., 2021). Although suggestive, the hypothesis of *Bdellovibrio* associated with *L. venatrix* was not proven because of negative FISH results. Furthermore, we could not find bacteria with that peculiar morphology, i.e., comma-shaped and monoflagellated bacteria, in TEM sections. In general, the bacteria observed in the cytoplasm of TEM-processed *L. venatrix* cells were not flagellated; however, it is well known that flagella can be lost or overlooked in TEM processed material due to their fragile structure and low (to very low) thickness. In this regard, the OTUs #2 and #4 (Table 2) seemed to represent environmental contaminants based on 1. *Pseudomonas composti* and *Panacagrimonas perspica* easily found as free-living bacteria and showing rod-shaped cells equipped with flagella (Im et al., 2010; Gibello et al., 2011) not detected in our ultrastructural analyses, and 2. the lack of FISH positive signals. As for OTU #3 (Table 2), *Agrobacterium tumefaciens,* it is a flagellated, bacilloid bacterium commonly associated with soil or plants (Escobar & Dandekar, 2003). Unfortunately, our FISH results did not help to identify the bacteria observed in the cytoplasm of *L. venatrix*; however, it should be noted that, according to our TEM analysis, bacteria were not abundant in the ciliate cytoplasm, which could have significantly affected FISH output.

Regarding the OTUs #5–11 (Table 2) found in *P. famelica*, we should be cautious about ascribing any of them to the status of symbionts, as we did not have the opportunity to perform any FISH experiments. Nevertheless, staying in the realm of speculation, we can hypothesize that the most likely candidate for a symbiont role could be OTU #11 - *Cysteiniphilum* sp., given the higher coverage value and that this genus is closely related to the pathogenic bacterium *Francisella tularensis* (Qian et al., 2023) (*Gammaprotobacteria*) which has been already found in association with ciliates, such as the cases of “*Candidatus* Francisella noatunensis subsp. endociliophora” in *Euplotes raikovi* (Schrallhammer et al., 2011) and *F. adeliensis* described from *E. petzi* (Vallesi et al., 2019). Considering the phenotypic characteristics of *Cysteiniphilum* (and the *Francisellaceae* as well) including that they are gram-negative, nonmotile, coccobacilli-shaped bacteria, at least two of the four types of bacteria discovered in the cytoplasm of *P. famelica* by means of TEM might be putatively interpreted as *Cysteiniphilum,* i.e., the homogeneously electrondense, irregularly shaped bacteria and the almond-shaped bacteria, with a lighter cytoplasm (Fig. 6A, C). Both the latter types of bacteria are in direct contact with the *Phialina*’s cytoplasm and encircled by a white halo, which is a common trait with “*Candidatus* Francisella noatunensis subsp. endociliophora” in *Euplotes raikovi* (Schrallhammer et al., 2011). Additionally, the more electrondense bacteria were especially observed around *Phialina*’s macronucleus, recalling the situation of *F. adeliensis* described from *E. petzi* (Vallesi et al., 2019). Unfortunately, as mentioned earlier, we cannot currently confirm morphological and molecular data using FISH data. Therefore, while it is important to describe ciliates and their associated microbial consortia to improve species descriptions (Serra et al. 2020), we are currently engaging in mere speculation.

In *C. vermicularis* we detected a single 16S rDNA sequence (OTU #12, Table 2) closely associated with “*Ca.* Amoebophilus”, common endosymbionts of *Acanthamoeba*, such as “*Ca.* Amoebophilus asiaticus” (Horn et al., 2001; Choi et al., 2009). Indeed, our TEM images showed rod-shaped bacteria similar in size and morphology to those previously observed in *Acanthamoeba* (Horn et al., 2001): they share a clear non-homogeneous cytoplasm and a double membrane plus the association, often, with many host’s ribosomes. However, the amoeba’s bacterium is encircled by a symbiosome bearing the ribosomes which is considered hypothetically derived from the rough endoplasmic reticulum, whereas in our case expanded rough endoplasmic reticulum cisternae, constituting a feature shared by all the three *Chaenea* species investigated so far for their ultrastructure (*C. vorax*, *C. teres*, *C. vermicularis*), are frequently associated with not symbiosome-included bacteria. This observation suggests the occurrence of a high exchange of metabolites between *C. vermicularis* and the bacteria, which surely deserves further investigation. Additionally, in the amoeba there are clusters of endobacteria which were not observed in *C. vermicularis*. Given the genetic distance with the sequence deposited in NCBI, we can hypothesize that the organism recovered in *C. vermicularis* likely belongs to another species of the same genus or another family respect to “*Ca.* Amoebophilus asiaticus” that has yet to be described.

As a final remark, some haptorians (e.g., *Arcuospathidium, Chaenea*, *Lacrymaria*) appear to be adapted to anaerobic habitats by harboring endosymbiotic methanogens (Finlay & Maberly, 2000). While ectosymbionts have been described on the surface of *C. teres* occasionally (Lipscomb and Riordan, 1990), we did not observe any kind of ectosymbionts on the surface of *L. venatrix*, *P. famelica,* and *C. vermicularis* (Figs. 7, 8A, D, E, 9).

## CONCLUSIONS

This study provides a comprehensive morphological and ultrastructural characterization, combined with molecular analyses, of three novel litostomatean ciliates: *Lacrymaria venatrix* sp. nov., *Phialina famelica* sp. nov., and *Chaenea vermicularis* sp. nov. The identification and distinction of these species from their respective congeners were achieved through an integrative approach that included live observation, silver impregnation, electron microscopy, and 18S rDNA gene-based phylogenetic analysis—highlighting the crucial role of integrative taxonomy in elucidating ciliate biodiversity.

In addition, we sequenced and annotated the mitogenomes of these species, which are among the few currently available in online databases, and analyzed their associated microbial consortia. Beyond the detailed species descriptions, we also address the questionable taxonomic status of *Lacrymaria* and *Phialina*, whose phylogenetic positions reveal a non-monophyletic relationship, as well as that of *Chaenea*, whose clade appears notably divergent from the other members of Haptoria.

## NOMENCLATURAL ACTS

The present work has been registered in ZooBank (code: urn:lsid:zoobank.org:pub:80925A7B-C1FB-4771-A070-46D3B4130095), as well as *Lacrymaria venatrix* sp. nov. (code: urn:lsid:zoobank.org:act:37BAE6A5-8A25-4BA0-84FA-E722C06083F9), *Phialina famelica* sp. nov. (code: urn:lsid:zoobank.org:act:3E3E5D0F-1D75-436F-B107-C2CEA3999B45), and *Chaenea vermicularis* sp. nov. (code: urn:lsid:zoobank.org:act:B0CD817E-B0EA-438F-A5C9-DF6C0AD4EBBE). The correspondent web pages are available at the following addresses: https://www.zoobank.org/References/80925A7B-C1FB-4771-A070-46D3B4130095, https://www.zoobank.org/NomenclaturalActs/37bae6a5-8a25-4ba0-84fa-e722c06083f9, https://www.zoobank.org/NomenclaturalActs/3E3E5D0F-1D75-436F-B107-C2CEA3999B45, and https://www.zoobank.org/NomenclaturalActs/b0cd817e-b0ea-438f-a5c9-df6c0ad4ebbe, respectively.

## DATA AVAILABILITY STATEMENT

The data that support the findings of this study are openly available in free access on-line repository or are available from the authors upon reasonable request.

## ACKNOWLEDGEMENTS

Special acknowledgements to Simone Gabrielli for photo-editing support. This work was supported by the University of Pisa PRA_2018_63 project. The research leading to these results has received funding from the European Community’s H2020 Programme H2020-MSCA-RISE 2019 under grant agreement n° 872767. The present study was also supported by the Fondazione Cassa di Risparmio di Pistoia e Pescia - giovani@ricercascientifica2019 fellowship (n° 2019.0380).

## CONFLICTS OF INTEREST

Authors declare that they do not have any conflict of interest.

## Notes

### Competing Interest Statement

The authors have declared no competing interest.

