## Supplementary Table 1 for "Description of three new predator species of Litostomatea (Alveolata, Ciliophora), including sequencing and annotation of their mitochondrial genomes"

| **Supplementary Table 1.** **List of known proteins found in the three litostomatean analyzed genomes**. List of proteins, in the three mitochondrial genomes, for which was possible determine the function. | | | |
| --- | --- | --- | --- |
| **Gene** | ***Lacrymaria venatrix*** | ***Phialina famelica*** | ***Chenea vermicularis*** |
| *atp9* | present | absent | present |
| *cob* | present | present | present |
| *cox1* | present | present | present |
| *cox2* | present | present | present |
| *nadh1_a* | present | absent | absent |
| *nadh10* | present | present | absent |
| *nadh2_b* | absent | present | present |
| *nadh3* | absent | absent | present |
| *nadh4L* | present | absent | absent |
| *nadh4* | present | present | absent |
| *nadh5* | present | present | present |
| *nadh7* | present | present | present |
| *nadh9* | present | present | absent |
| *rpL16* | present | present | absent |
| *rpL2* | absent | present | present |
| *rpL6* | absent | present | present |
| *rpS12* | present | present | present |
| *rpS13* | present | absent | absent |
| *rpS14* | present | present | absent |
| *rpS19* | absent | present | present |
| *rpS3_b* | present | present | absent |
| *rpS4* | present | absent | absent |
| *rpS8* | present | absent | absent |
| *rrnL* | present | present | present |
| *rrnS* | present | present | present |
| *tRNA-Phe* | present | present | present |
| *tRNA-Tyr* | present | present | present |
| *tRNA-Trp* | present | present | present |
